# Microarrayed human bone marrow organoids for modeling blood stem cell dynamics

**DOI:** 10.1101/2021.05.26.445803

**Authors:** Sonja Giger, Moritz Hofer, Marijana Miljkovic-Licina, Sylke Hoehnel, Nathalie Brandenberg, Romain Guiet, Martin Ehrbar, Esther Kleiner, Katharina Gegenschatz, Thomas Matthes, Matthias P. Lutolf

## Abstract

In many leukemia patients, a poor prognosis is attributed either to the development of chemotherapy resistance by leukemic stem cells (LSCs) or to the inefficient engraftment of transplanted hematopoietic stem/progenitor cells (HSPCs) into the bone marrow (BM). Here, we build a 3D in vitro model system of bone marrow organoids (BMOs) that recapitulate several structural and cellular components of native BM. These organoids are formed in a high-throughput manner from the aggregation of endothelial and mesenchymal cells within hydrogel microwells. Accordingly, the mesenchymal compartment shows partial maintenance of its self-renewal and multilineage potential, while endothelial cells self-organize into an interconnected vessel-like network. Intriguingly, such a vascular compartment enhances the recruitment of HSPCs in a chemokine ligand/receptor-dependent manner, reminiscent of HSPC homing behavior *in vivo*. Additionally, we also model LSC migration and nesting in BMOs, thus highlighting the potential of this system as a well accessible and scalable preclinical model for candidate drug screening and patient-specific assays.

## Introduction

Bone marrow (BM) governs hematopoietic stem/progenitor cell (HSPC) homeostasis and differentiation through its complex microenvironment^1^, which comprises two distinct HSPC niches: one in the microvessel-enriched trabecular region, and another at the cortical regions of the bones^2^. Within these niches, HSPC behavior is tightly regulated by a multitude of cellular players, including mesenchymal stem/progenitor cells (MSCs), vascular endothelial cells (ECs), osteoblasts, sympathetic nerve fibers, macrophages, megakaryocytes, adipocytes, and non-myelinating Schwann cells^3,4^. However, despite extensive control mechanisms, a variety of hematological and immunological diseases still occur, including acute myeloid leukemia (AML), with its defining characteristic attributed to the rapid accumulation of abnormal immature progenitor cells^5,6^. Classically, the treatment of AML consists of chemotherapy followed by transplantation of healthy HSPCs, yet this often fails to substantially improve patient prognosis^7^. Relapses are mainly attributed either to the inability of the treatments to eliminate all leukemic blasts or the failure of injected donor HSPCs to engraft into the patient’s BM; both failures are associated with a misfunctioning BM niche microenvironment. Thus, modeling human BM with its intricate stem cell-niche interactions is pivotal for the development of novel therapeutic strategies.

Classic *in vivo* models (e.g., xenograft mice) used to assess the homing capacity, engraftment, and reconstitution potential of donor HSPCs have significantly advanced the general understanding of HSPC behavior^8^, though they remain tightly linked to ethical and technical constraints^9^. Bioengineered cell culture systems have been proposed as potential alternatives for the manipulation and observation of HSPC behavior, such as self-renewal and differentiation in niche-mimicking microenvironments^10–12^. However, these *in vitro* systems are based on two-dimensional (2D) culture settings and, as such, poorly recapitulate the complex and multicellular three-dimensional (3D) environment found in the native BM.

Stem cell-derived self-organizing structures, so-called organoids, have recently emerged as a revolutionary tool for basic and translational research^13,14^. These multicellular *in vitro* structures recapitulate the cellular spatio-temporal organization that occurs during development, tissue homeostasis, and regenerative processes *in vivo*^15^. However, despite extraordinary activity in the organoid field over the past decade, very few *in vitro* organoid model systems for BM have been described. The closest approximations are likely those developed through bioengineering techniques, such as “ossicle” scaffolds^16,17^, which are based on the *ex vivo* formation of a human stromal cell layer followed by a subsequent ectopic transplantation. Such engineered ossicles show comparable morphological, phenotypic, and functional features to native bone organs, facilitating the probing of “humanized” microenvironments for human HSPCs *in vivo*^16,17^. However, the dependency of this system on xenotransplantation remains a major drawback for standardization. A fully *in vitro* setup has also been established, using a perfusion bioreactor system to form an engineered human osteoblastic niche^18^. Even more relevantly, the formation of scaffold-free mesenchymal spheres, called mesenspheres, has been shown to preserve the multilineage potential of HSPCs by secreting HSPC-supportive cytokines^19^. Despite mimicking the phenotypic and structural characteristics of the BM mesenchymal compartment, both of these systems still fail to recapitulate the multicellular context of the native BM. Two recent studies integrated such multicellular dimensions into “bone marrow-on-chip” platforms, built by a microfluidic channel containing stromal cells that is positioned parallel to a channel lined with endothelial cells^20,21^. Partial maintenance of HSPCs and their multilineage differentiation has been observed in these systems, yet their wider applicability may be limited by the static and indirect interaction of the two cell types as well as the low scalability of the designs^20,21^.

To close this technology gap, we have developed a novel approach for generating BM organoids (BMOs) with *in vivo*-like functional characteristics. Here, we see the self-organization of a pseudo-vascularized network within a mesenchymal compartment via the aggregation of the two major niche cell types, MSCs and ECs. Intriguingly, BMOs containing such a vascular compartment displayed an enhanced recruitment of HSPCs, which then resided in close proximity to the endothelial network. Furthermore, this migratory behavior seems to be partially regulated in a chemokine ligand/receptor-dependent manner, similar to the *in vivo* process. We also provide proof-of-concept for the deployment of this system for disease modeling, particularly in AML. We thus believe that this organoid system holds great promise as a preclinical model for pharmacokinetic assays and toxicity studies of novel drugs in a BM niche-mimicking 3D environment.

## Results

### Formation of bone marrow organoids in a scalable manner

Guided by the natural composition of BM, we selected two major cellular niche components, namely MSCs and ECs, for the generation of BMOs. We used a commercially available microwell platform consisting of a high-density array of round-bottom poly(ethylene glycol) (PEG) hydrogel microwells (Gri3D^®^) to generate BMOs in a high-throughput manner (**Fig. 1a**)^22^. Once sedimented at the bottom of the microwells, the two cell types formed a compact mesenchymal spheroid featuring a morphologically distinct vascular compartment (**Fig. 1a**). Considering the essential role of MSCs and ECs in the recruitment and maintenance of HSPCs^23,24^, we hypothesized that these cells could trigger the homing and engraftment of HSPCs and leukemic blasts in BMOs, thus mimicking organ-level physiological processes (**Fig. 1a**).

**Fig. 1.**
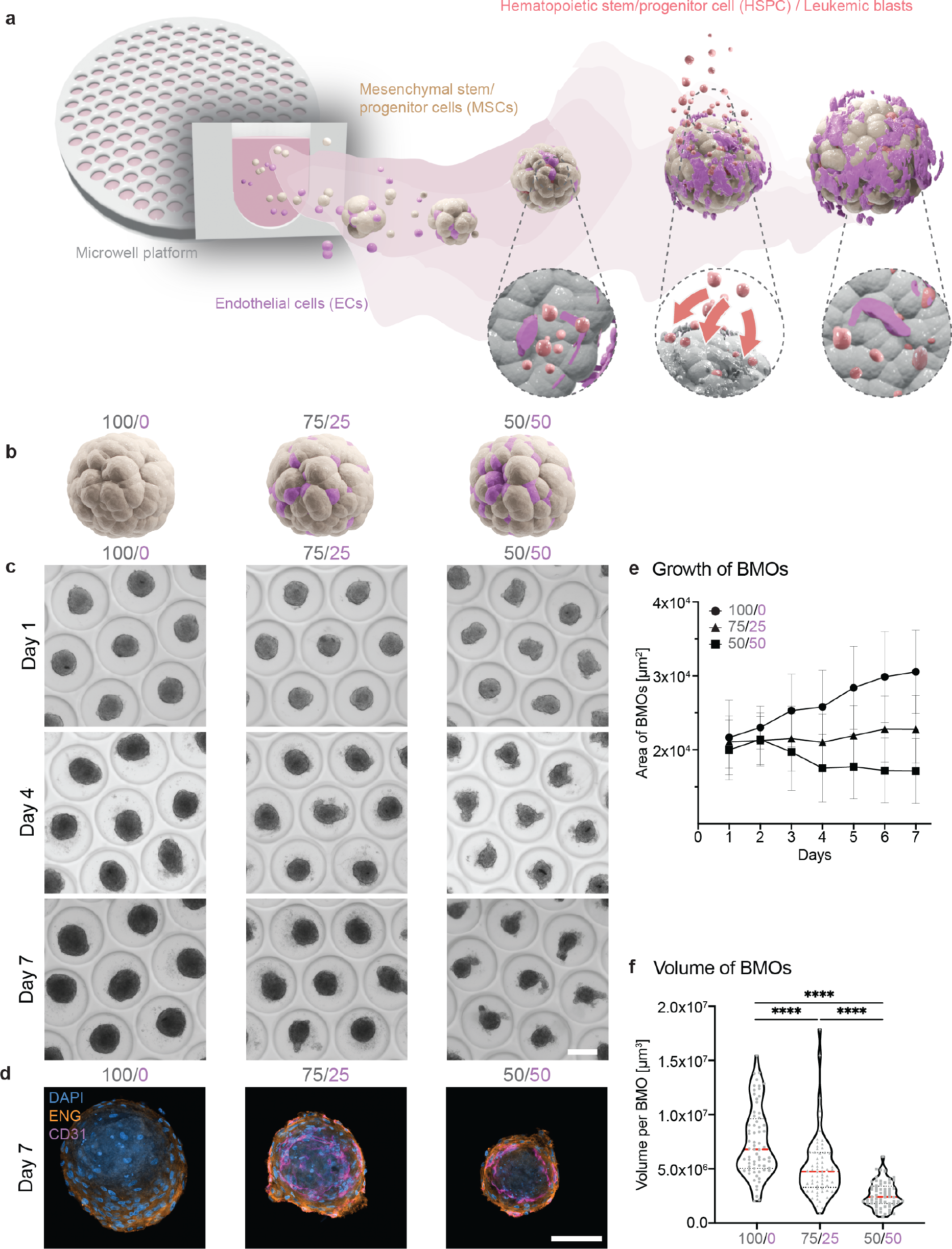
Formation of bone marrow organoids in a scalable manner. **a**) Schematic illustration of the concept of BMOs. Self-aggregation of MSCs (beige) and ECs (magenta) in the microwell platform for the formation of BMOs. After the initial aggregation, HUVECs self-organize to form a vascularization-like compartment within the mesenchymal tissue. HSPCs and leukemic blasts (red) are attracted by the presence of multiple BM cell types in a homing-like behavior and take up residence in this BM-mimicking environment. Together with the easily accessible and scalable characteristics of BMOs, they offer a promising in vitro 3D model system for BM research and drug candidate screening. **b**) Schematic illustration of the aggregation of different ratios of MSCs (beige) and ECs (magenta). The control condition consists of only mesenchymal cells (100/0, left) and the two multi-cell type conditions contain either 25 % (75/25, center) or 50 % (50/50, right) endothelial cells. **c**) Representative brightfield images of BMOs with different cell ratios in the Gri3D platform at different time points (Day 1, 4, 7). Scale bar, 200 μm. **d**) Representative confocal images of the mesenchymal marker expression endoglin (ENG, orange), the endothelial marker CD31 (magenta), and DAPI (blue) marking the cell nuclei in BMOs. Scale bar, 100 μm. **e**) Growth of BMOs via the analysis of brightfield images by an automated script in Fiji. Analysis of 300–1,600 BMOs per condition. Symbols indicate the mean values, and the error bars show the standard deviation. Results of four independent experiments. **f**) Spheroids were cultured for 7 days, and after fixation, the estimated BMO volume was calculated based on a mask made from the data of ENG expression on the surface of BMOs. Violin plots represent the smoothened distributions, red horizontal lines indicate the medians, and black dotted lines the quartiles. Results of four independent experiments. Statistical analysis by ordinary one-way ANOVA with multiple comparisons. ****p < 0.0001.

To define the optimal 3D culture condition for the formation of BMOs, we first tested the aggregation capacity of human MSCs and human umbilical vein endothelial cells (HUVECs) in two different ratios, namely 50/50 and 75/25, respectively, compared to a 100 % MSCs control condition (100/0)^19^ (**Fig. 1b**). Bright-field imaging demonstrated an efficient and robust generation of almost 3,000 BMOs per Gri3D^®^ 24-well plate (121 BMOs/well) (**Fig. 1c; Supp. Fig. 1a**). After 7 days in culture, all tissues stained positively for the mesenchymal marker endoglin (ENG) and the endothelial marker CD31 for dual-cell type conditions (50/50; 75/25) (**Fig. 1d**); HUVECs seeded alone in the microwells neither aggregated nor survived (data not shown). The largest aggregate growth was detected in the 100/0 control condition followed by the 75/25 condition, and the smallest growth was detected for the 50/50 condition (**Fig. 1e**). Increasing the percentage of ECs resulted in smaller, yet more reproducible, aggregate sizes, which was also confirmed by quantitative volumetric analyses (**Fig. 1f**). These data demonstrate that our high-throughput culture method offers a robust and efficient formation of large quantities of BMOs composed of MSCs and ECs.

### Self-renewing and differentiated MSCs in bone marrow organoids

We next characterized the composition and identity of the mesenchymal compartment within BMOs by assessing the presence of key phenotypic markers. Immunofluorescence analysis revealed comparable expression levels among all conditions for nestin (NES) and transgelin (TAGLN), two mesenchymal factors involved in HSC recruitment and blood vessel formation *in vivo*^19,25^ (**Fig. 2a,b; Supp. Fig. 2a**). Interestingly, we found a higher expression of the self-renewal mesenchymal marker ENG in the dual-cell type BMOs (75/25; 50/50) compared to the control condition (100/0) (**Fig. 2c; Supp. Fig. 2a**). To quantitatively assess the portion of multipotent, self-renewing MSCs characterized by the CD71^−^/CD31^−^/CD45^−^/ENG^+^/CD146^+^ immunophenotype^19,24^, we analyzed digested BMO cells by flow cytometry (**Supp. Fig. 2b,c**). Within the CD71^−^/CD31^−^/CD45^−^ mesenchymal cells, we observed a higher mean fluorescent intensity (MFI) for ENG in the dual-cell type BMOs, supporting our immunofluorescence observations, while the CD146 levels remained constant (**Supp. Fig. 2d**). In terms of cell numbers, we detected an increase of up to 30–50 % of the ENG^+^/CD146^+^ double-positive (self-renewing) phenotype in both dual-cell type conditions compared to the mesenchymal control condition (**Fig. 2d; Supp. Fig. 2e**). These findings show that the phenotypic self-renewing MSCs are more abundant in the dual-cell type bone marrow organoids, possibly through a supportive interaction between the ECs and MSCs, compared to HSPC recruitment or blood vessel formation.

**Fig. 2.**
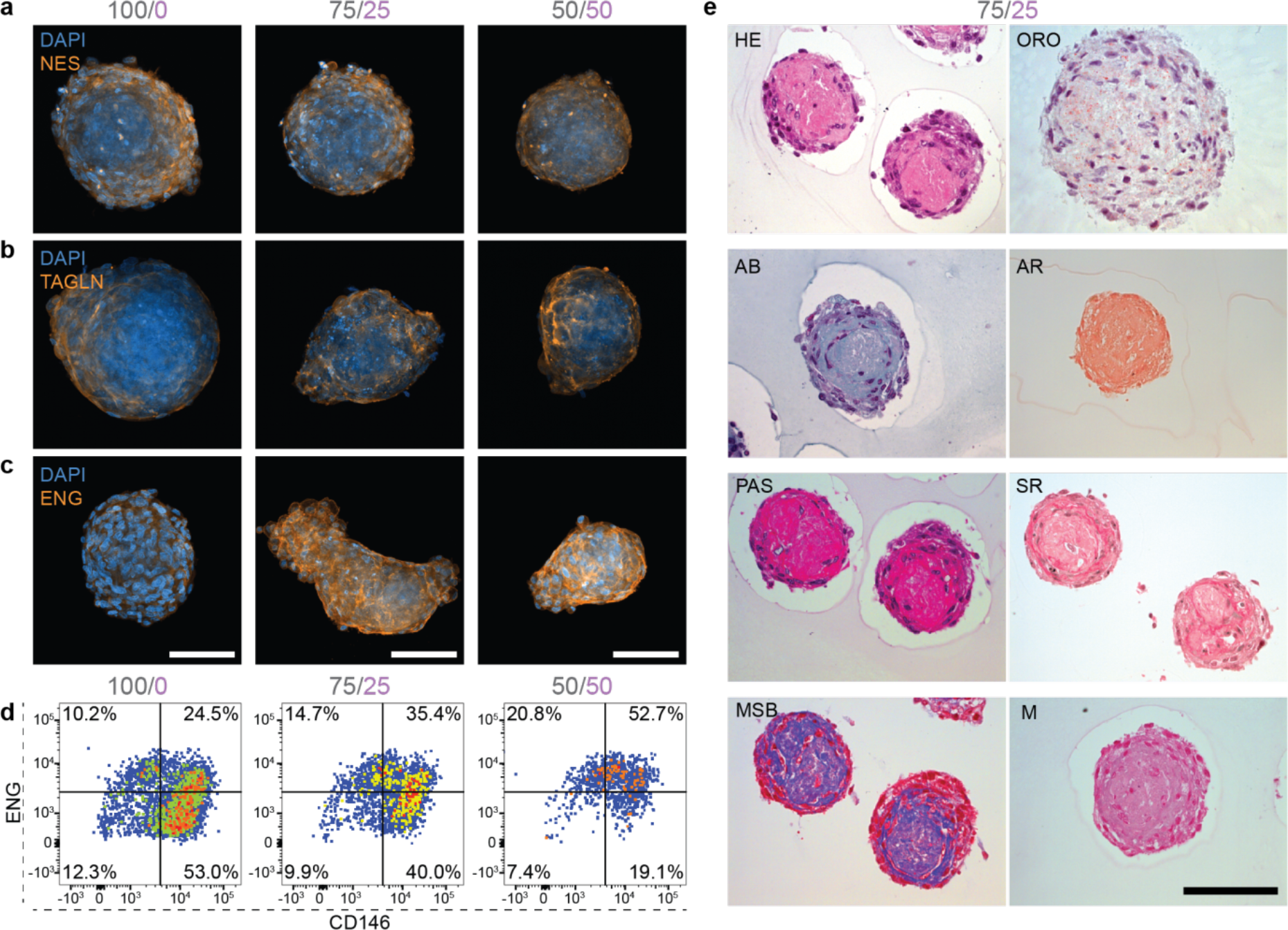
Self-renewing and differentiated MSCs in bone marrow organoids. **a, b, c**) Representative immunofluorescence images of the expression of mesenchymal markers (**a**) nestin (NES, orange), (**b**) transgelin (TAGLN, orange), and (**c**) endoglin (ENG, orange) in BMOs cultured for 7 days. Maximum intensity projection of z-stack confocal images. Scale bars, 100 μm. **d**) Representative flow diagrams show the percentage of ENG+ and CD146+ cells of the CD71-/CD31-/CD45-population in the three different conditions (100/0, 75/25, 50/50). **e**) Representative images of histological staining of the 75/25 BMO condition cultured for 7 days. The morphology is represented by hematoxylin and eosin (HE) staining and Oil Red O (ORO)-stained lipid droplets. The Alcian blue (AB) staining marks acidic polysaccharides, such as glycosaminoglycans, and alizarin red (AR) detects calcium deposits. Periodic Acid Schiff (PAS) stains for polysaccharides, Sirius red (SR) stains for collagen, Marius Scarlett Blue (MSB) stains for fibrin, and Miller (M) stains for elastin. Scale bar, 100 μm.

Using histology, we next asked if MSC-derived differentiated cell types appeared in BMOs. Hematoxylin and eosin (HE) staining showed that the majority of the cells grow at the periphery of the structures, while the center appears as a dense eosinophilic core (light pink, **Fig. 2e**). The eosinophilic core had a low expression of dead cell markers (data not shown), suggesting that an extracellular matrix is likely secreted in the center of the BMOs. Further histological analyses revealed an abundance of collagen, fibrin, and polysaccharides, such as glycosaminoglycans, which are known extracellular matrix components in the MSCs residency site in natural BM^26^; however, there was no elastin accumulation (**Fig. 2e**). Furthermore, these deposits suggest the commitment of lineage-specific progenitors, such as chondrocytes, osteocytes, and adipocytes in BMOs. The presence of the latter is supported by Oil Red O (ORO) staining that revealed the existence of lipid–droplet-bearing adipocytes^27^, indicating that a subset of MSCs underwent adipogenic differentiation (**Fig. 2e**). Next, the detection of SRY-box transcription factor 9 (SOX9), a marker required for the onset of chondrogenesis, supports the partial commitment towards the chondrogenic lineage (**Supp. Fig. 2f**)^26,28^. Additionally, the Runt domain-containing transcription factor 2 (RUNX2), a transcription factor regulating osteogenic MSC differentiation^26^, was detected in the co-cultures (75/25; 50/50) (**Supp. Fig. 2g**). However, no calcium deposits were detected after the 7-day cultivation period in BMOs, indicated by negative alizarin red (AR) staining (**Fig. 2e**). Overall, this phenotypic analysis shows that the co-culture of MSCs and ECs in BMOs positively influences the maintenance of self-renewing MSCs and simultaneously allows a partial maturation into adipocytes, chondrocytes, and pre-osteoblasts.

### Establishment of a self-organized, vascular-like network

We further characterized the capacity of the endothelial cells to form vascular-like structures within the mesenchymal cells by immunofluorescence analysis. In contrast to aggregates consisting purely of mesenchymal cells, those containing endothelial cells displayed a network structure, evidenced by the general endothelial markers CD31 and CD144 (**Fig. 3a; Supp. Fig. 3a,b**). Notably, higher numbers of ECs in the initial culture condition did not result in an increased proportion of CD31^+^-expressing cells (**Fig. 3b; Supp. Fig. 3c,d**), suggesting that EC survival might depend on the relative number of MSCs in the co-culture. The expression of the endothelial-specific adhesion molecule CD144 within the CD31^+^ population further suggests that endothelial cell-cell contacts are established through surface junctions (**Supp. Fig. 3e**), whereas active sprouting seems to occur very rarely, indicated by low vascular endothelial growth factor receptor (VEGFR) expression (**Supp. Fig. 3f**). These results demonstrate that ECs can self-organize into an interconnected endothelial network in a 3D co-culture with MSCs.

**Fig. 3.**
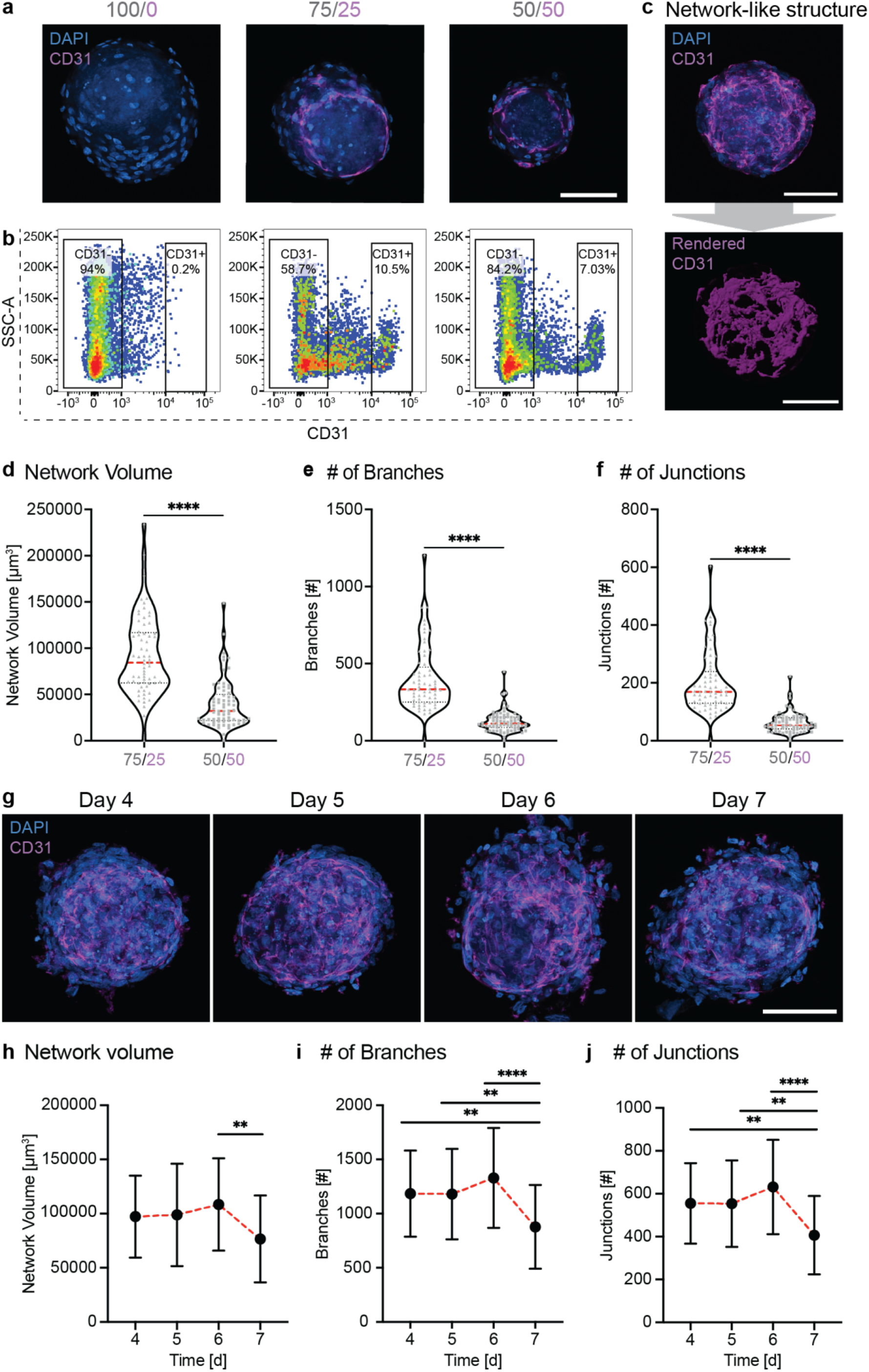
Establishment of a self-organized, vascular-like network. **a**) Representative single-plane confocal images of the endothelial marker CD31 (magenta) and DAPI (blue) in the different BMOs (100/0, 75/25, 50/50). Scale bar, 100 μm. **b**) Representative flow diagrams show the percentage of CD31+ cells among all live cells of BMOs from the different conditions (100/0, 75/25, 50/50). **c**) Representative maximum intensity projection of a z-stack image of CD31 (top panel) and the resulting rendered data (bottom panel) of the self-organized vascular network-like structure in the BMO condition 75/25. Scale bar, 100 μm. **d, e, f**) Quantification of the network architecture by estimating (**d**) the volume, (**e**) the number of branches, and (**f**) junctions. Results from four independent experiments. Statistical analysis by Student’s t-test. ****p < 0.0001. **g**) Representative images of the endothelial marker CD31 in BMOs at day 4, 5, 6, 7. Maximum intensity projection of z-stack confocal images. Scale bar, 100 μm. **h, i, j**) Quantification of the endothelial network (**h**) volume, (**i**) branches, and (**j**) junctions at different time points (day 4, 5, 6, 7). Results of three independent experiments. Symbols indicate mean values and the error bars show the standard deviations. Statistical analysis by ordinary one-way ANOVA and multiple comparisons. **p < 0.01; ****p < 0.0001.

The complexity of such endothelial network architecture was visualized and quantified by 3D-rendered and skeletonized data based on CD31 immunofluorescence images (**Fig. 3c; Supp. Fig. 3g**). This quantification showed that the largest and most complex networks, in terms of number of junctions and branches, were formed in the 75/25 condition (**Fig. 3d-f**). Subsequently, we analyzed the evolution of the network architecture in this condition over 7 days of culture (**Fig. 3g**). We measured stable network parameters for 6 days, with a slight drop in both network volume and the number of branches and junctions on the seventh day in culture (**Fig. 3h-j**). Altogether, our finding of a stable, interconnected, vascular-like network in dual-cell type bone marrow organoids suggests that this 3D co-culture system partially resembles the vascular niches within the native BM environment.

### Bone marrow organoids as a 3D migration assay

Next, we aimed at developing a 3D-*in vitro* assay with these BMOs to investigate the homing ability of human HSPCs (**Fig. 4a**). To that aim, HSPCs derived from cord blood samples (see methods, **Supp. Fig. 4a**) were added to three-day-old BMOs with or without a vascular-like network (**Fig. 4a**). After an additional 4 days of culture, the number of successfully homed HSPCs within the BMOs was analyzed by immunofluorescence staining (**Fig. 4a**). This analysis demonstrated the presence of homed HSPCs marked by the pan-hematopoietic marker CD45 within BMOs in all three conditions (**Fig. 4b; Supp. Fig. 4b**). Strikingly, BMOs containing ECs displayed a significantly higher number of homed CD45^+^ HSPCs compared to aggregates consisting purely of mesenchymal cells (**Fig. 4c**). This result emphasizes the importance of the vascular compartment in BMOs for modeling HSPC homing behavior *in vitro*. Titration experiments showed that the number of successfully homed HSPCs was dependent on the number of initially seeded cells (**Fig. 4d**). To further characterize the homing behavior, we analyzed the location of the HSPCs within the BMOs by quantifying their relative distance from the surface of the organoid (D_o_) and from the network (D_n_) based on the 3D-rendered data (**Fig. 4e**). Generally, the relative frequency distribution showed that HSPCs migrate, on average, deeper into BMOs than what would be expected when assuming a uniform distribution within the organoid volume (25–30 μm, black dotted line) (**Fig. 4f**). The observation that ECs are preferentially arranged on the outer region of BMOs could explain why HSPCs in the dual-cell type BMOs typically reside closer to the organoid surface (**Fig. 4f**). This observation was confirmed in both dual-cell type conditions, where over 90 % of HSPCs were closer to the endothelial network structure when compared to the distance expected when generated from a modeled random distribution (9.8 ± 4.1 μm in 75/25, and 10.0 ± 6.8 μm in 50/50) (**Fig. 4g**). First, these results show that HSPCs can efficiently migrate inside the organoid structure, and second, the abundance of ECs in BMOs positively influences the migratory behavior of HSPCs in this 3D-homing assay. Overall, BMOs containing the largest vascular-like network (75/25) resulted in the most efficient homing of HSPCs, suggesting that these BMOs provide an optimal environment for such a 3D-migration assay.

**Fig. 4.**
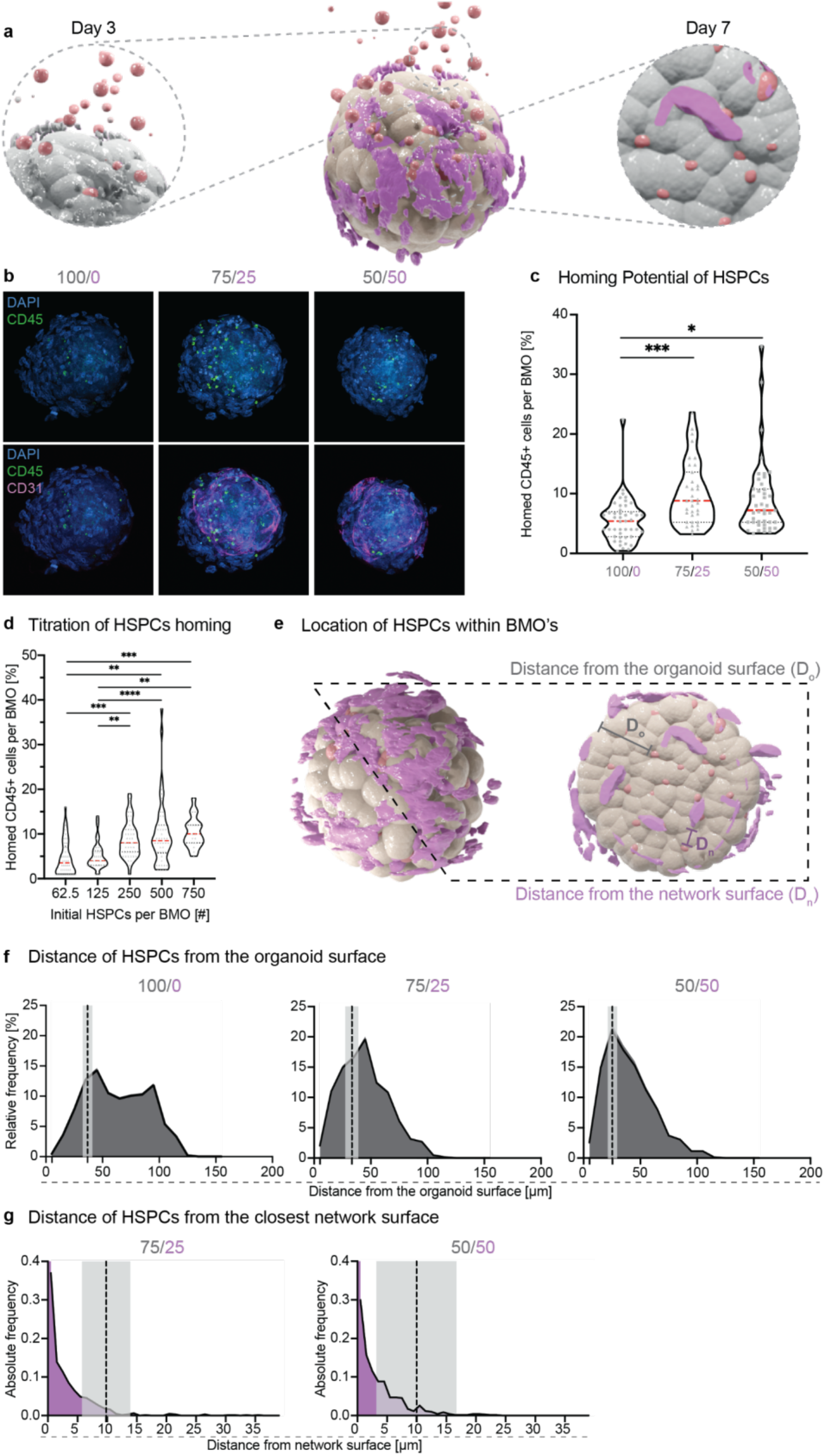
Bone marrow organoids as a 3D migration assay. **a**) Schematic representation of the seeding of HSPCs (red) onto BMOs on day 3 and the subsequent analysis of efficiently homed HSPCs on day 7. **b**) Representative confocal images of the HSPCs (CD45, green) and the network (CD31, magenta) in BMOs (DAPI, blue) from the three conditions (100/0, 75/25, 50/50). The maximum projection of z-stack acquisition is shown. Scale bar, 100 μm. **c**) Quantitative analysis of the percentage of homed CD45+ cells per BMO normalized to the initial number of seeded HSPCs. Violin plots represent the distributions, red horizontal lines indicate the medians, and black dotted lines the quartiles. Results of four independent experiments. Statistical analysis by ordinary one-way ANOVA and multiple comparisons. *p < 0.05; ***p < 0.001. **d**) Quantitative analysis of the number of homed CD45+ cells per BMO, derived from different initial seeding numbers of lineage-depleted HSPCs. Violin plots represent the smoothened distributions, red horizontal lines indicate the medians, and black dotted lines the quartiles. Results of three independent experiments. Statistical analysis by ordinary one-way ANOVA and multiple comparisons. **p < 0.01; ***p < 0.001; ****p < 0.0001. **e**) Schematic illustration of the definition of the distance to the organoid (Do) and network surface (Dn) in BMOs. **f**) Relative frequency distributions of the distance of the HSPCs from the organoid surface (Do) in the different BMO conditions (100/0, 75/25, 50/50). The dashed lines indicate the expected average distance by modeling per condition (100/0: 36.7 ± 4.2 μm, 75/25: 33.2 ± 5.6 μm, 50/50: 25.4 ± 4.7 μm; mean ± SD [shaded regions]). Around 79 % of HSPCs in the 100/0, 68 % in the 75/25, and 73 % in the 50/50 condition migrate deeper in BMOs than the expected distance. Results of three independent experiments. **g**) Absolute frequency distributions of the distance of the HSPCs from the endothelial network surface in the multicellular BMOs (75/25 and 50/50). The dashed lines indicate the average expected distance by modeling per condition (75/25: 9.8 ± 4.1 μm, 50/50: 10.0 ± 6.8 μm; mean ± SD [shaded region]). Around 94 % of HSPCs in the 75/25 and 92 % in the 50/50 condition reside closer to the endothelial structure in the BMO than the expected distance. Results of three independent experiments.

### Dynamics and mechanism of CD34^+^ HSPC homing

After showing that HSPCs have the potential to migrate inside the BMOs, we addressed the homing dynamics and mechanism of highly enriched HSPCs (CD34^+^ HSPCs, purity around 90 %) (**Fig. 5a; Supp. Fig. 5a**). The CD34^+^ HSPCs were pre-labelled with carboxyfluorescein succinimidyl ester (CFSE) prior to their addition to BMOs (see methods). Analogous to the previous procedure, 200 CFSE-labelled HSPCs were seeded on fully aggregated BMOs on day 3, and their occurrence within BMOs was analyzed over time by immunofluorescence analysis (**Fig. 5b; Supp. Fig. 5b**). The first CFSE^+^ cells were already detected within the BMOs 8 h after seeding (**Fig. 5c**). A constant increase of the number of CFSE^+^ cells was detected during the first 24 h, and a consistent, but slightly lower, increase was observed up until 96 h (**Fig. 5c**). This migratory behavior resembles *in vivo* HSPC homing behavior, which occurs around 20 h after intravenous injection into the BM of humanized mice^29^. Given that CD34^+^ stem cells are known to be a highly quiescent cell population^30,31^, the detected increase in cell number within BMOs is likely due to an increase in homing cells rather than proliferation. Indeed, a non-bimodal distribution of CFSE intensities over the time course supports this hypothesis (**Supp. Fig. 5c**), as CFSE intensity would decrease during cell division. These results also indicate that efficiently homed CFSE^+^ cells remain within the 3D niche for up to 4 days.

**Fig. 5.**
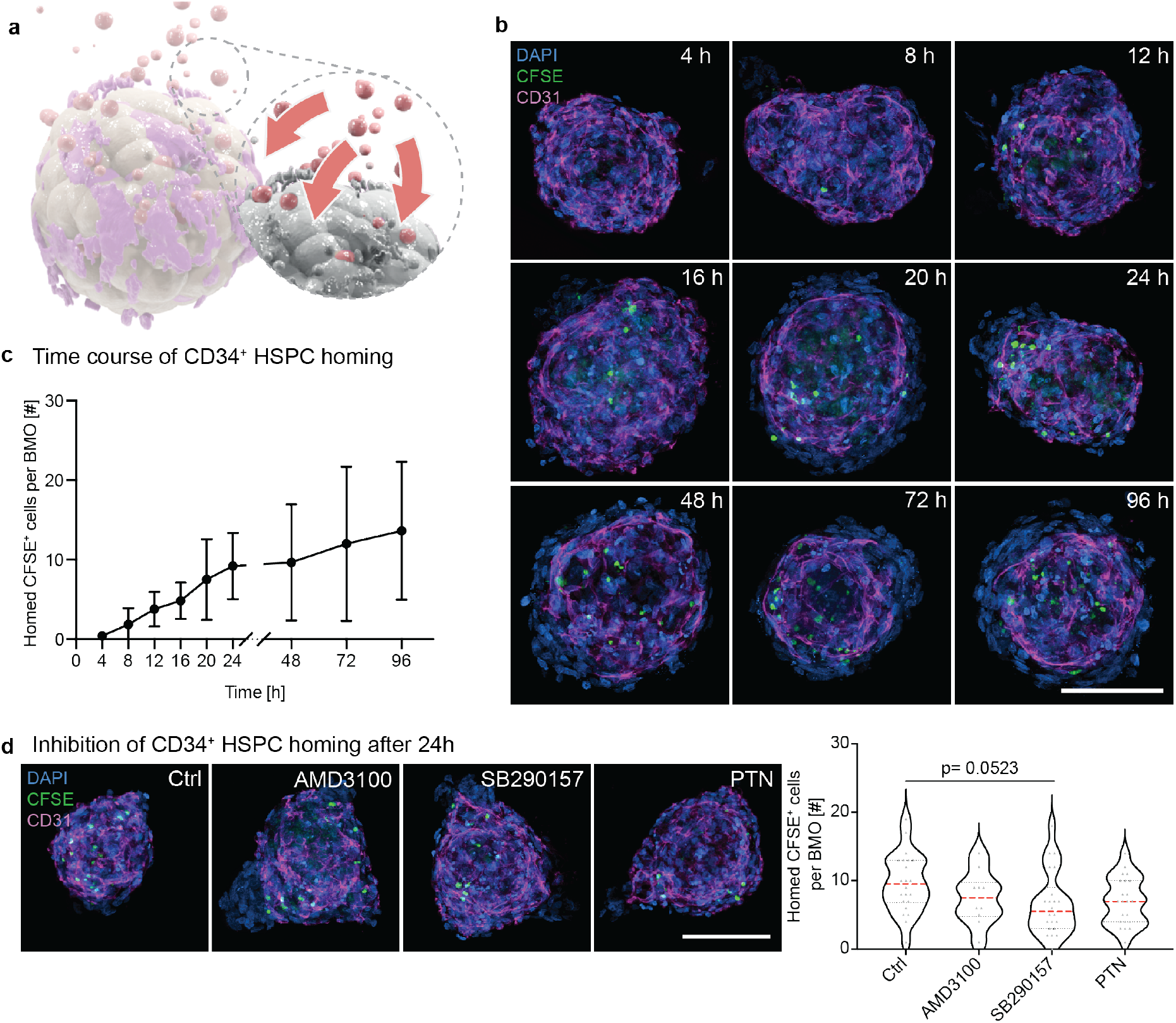
Dynamics and mechanism of CD34^+^ HSPC homing. **a**) Schematic illustration of the homing dynamics of HSPCs (red) into BMOs (magenta/beige/grey). **b**) Representative confocal images of CFSE-labeled HSPCs (CFSE, green) and the network (CD31, magenta) within BMOs (DAPI, blue) at 4 h, 8 h, 12 h, 16 h, 20 h, 24 h, 48 h, 72 h, and 96 h after seeding of HSPCs. The maximum projections of confocal z-stack acquisitions are shown. Scale bar, 100 μm. **c**) Quantitative analysis of the number of CFSE+ cells per BMO over time. Results of two independent experiments. Symbols indicate mean values, and the error bars show the standard deviations. **d**) Representative confocal images of the inhibition of homing of HSPCs into BMOs 24 h after seeding (left panel). The purified HSPCs (CFSE, green) are treated with either AMD3100 or SB290157 before seeding on BMOs. In the third condition, BMOs are treated with anti-PTN before seeding of the HSPCs to negatively influence the migration behavior of the HSPCs. The network is marked with CD31 (magenta) and the nuclei with DAPI (blue). The maximum projection of z-stack acquisition is shown. Scale bar, 100 μm. The number of homed CFSE+ cells is quantified per BMO per treatment (right panel). Violin plots represent the distributions, red horizontal lines indicate the medians, and black dotted lines the quartiles. Results of two independent experiments. Statistical analysis by ordinary one-way ANOVA and multiple comparisons.

Thereafter, we asked whether the homing of CD34^+^ HSPCs in BMOs is regulated via the C-X-C motif chemokine ligand 12 (CXCL12)/CXCR4 axis and/or the heparin-binding growth factor pleiotrophin (PTN)-signaling pathway. Both pathways have been found to be crucial for the lodgment, transmigration, and maintenance of HSPCs *in vivo*^32–34^. The chemokine CXCL12, which is secreted by bone marrow stromal cells, can attract CXCR4-expressing HSPCs to the BM niche^35,36^. Whereas PTN, which is mainly secreted by ECs, regulates the retention of HSCs in the BM niche via the interaction with its transmembrane receptor PTPRZ^33^. We found that CXCR4 was expressed by approximately 15 % of the CD34^+^ HSPCs (**Supp. Fig. 5d,e**) and PTN-positive cells were found broadly distributed throughout the BMOs (**Supp. Fig. 5f**). To test the involvement of CXCL12-CXCR4 interaction, we selected the CXCR4 antagonist AMD3100 and the C3aR antagonist SB290157, both known to negatively influence this interaction^37,38^. The first two conditions had CD34^+^ stem cells treated with either of these two inhibitors prior to seeding on BMOs, while the third condition included a pre-treatment of the BMOs with an anti-PTN antibody. All treatments showed a reduced number of CFSE^+^ cells within BMOs after 24 h, with the largest reduction observed in the SB290157 treatment (p = 0.0523) (**Fig. 5d**). These results suggest that the homing of HSPCs in BMOs is partly regulated by the CXCL12-CXCR4 signaling pathway, underlining the potential of the BMO system to model an *in vivo*-like homing behavior of HSPCs *in vitro*.

### Bone marrow organoids as a niche for leukemic blasts

Standard *in vitro* toxicology studies often fail to detect drug side effects, which can only be observed in elaborate and expensive *in vivo* studies. Moreover, the poor predictivity of drug responses from *in vitro* screenings have led to many unsuccessful clinical trials, due to the fact that cell responses differ based on whether they reside in their (patho-)physiological environment or an oversimplified *in vitro* setting. To showcase the potential value of BMOs for studying leukemia treatments, we tested the capacity of BMOs as a supportive environment for CD34^+^ leukemic blasts. Accordingly, leukemic blast cells were enriched for the CD34^+^ population and labelled with CFSE (**Supp. Fig. 6a**). This enriched population was then seeded on 3-day-old BMOs, and the number of internalized cells was assessed over time (**Fig. 6a; Supp. Fig. 5b**). Surprisingly, the first CFSE^+^ cells were detected within BMOs after only 4 h (**Fig. 6b**). The continuous increase of CFSE^+^ leukemic blast cells within the BMOs, together with a constant CFSE intensity over a 96 h timeframe, suggests an efficient migration of leukemic blasts rather than their proliferation within BMOs (**Fig. 6b,c**). Additionally, this rapid migration of leukemic blasts compared to healthy HSPCs correlates with a higher expression of CXCR4 on leukemic blasts (**Supp. Fig. 5e; Supp. Fig. 6c**). These results demonstrate that leukemic blasts can efficiently migrate in BMOs. Ultimately, BMOs inhabited by leukemic blasts could be used as an *in vitro* model system to allow large compound screenings on a multicellular entity.

**Fig. 6.**
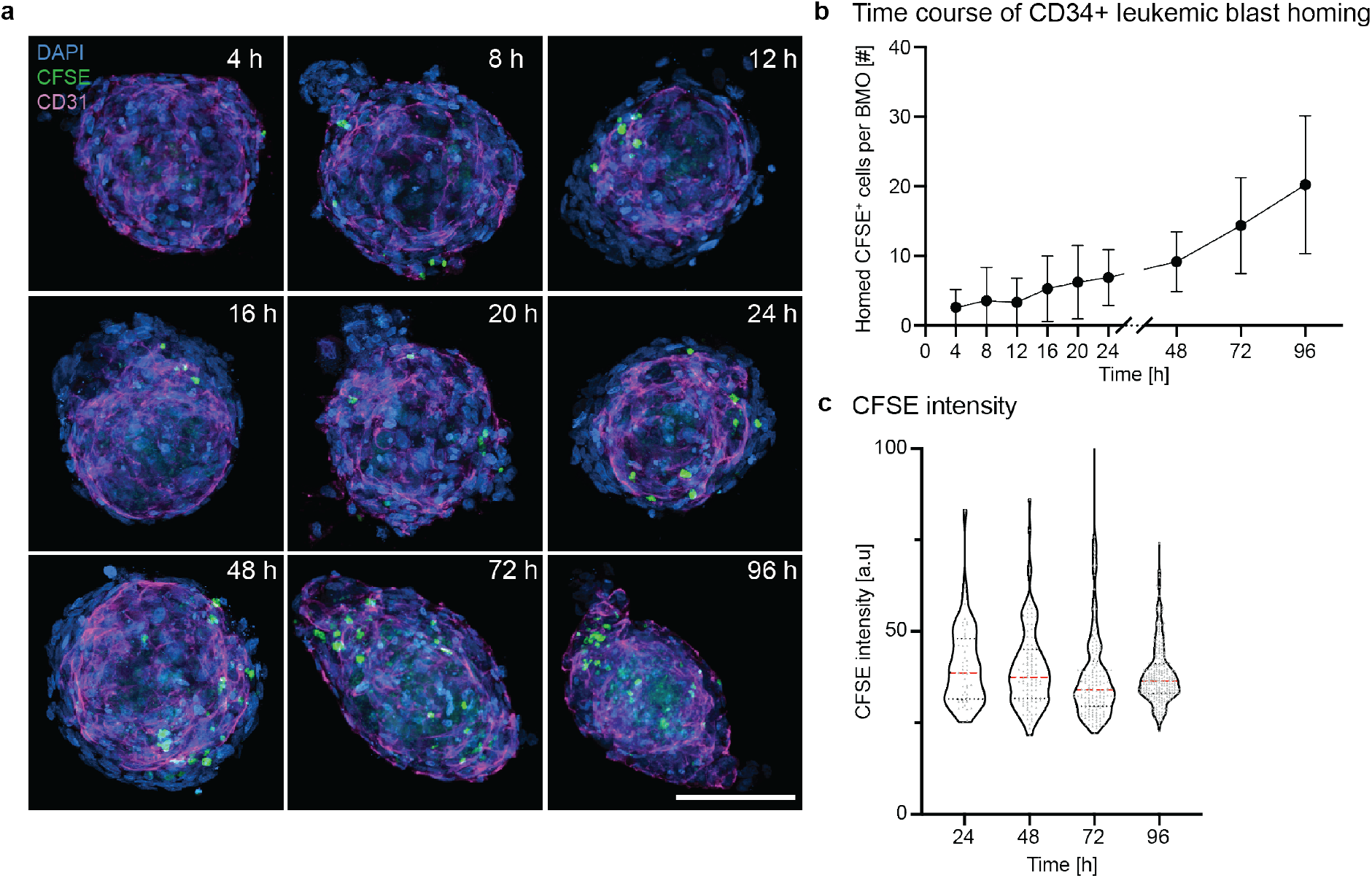
Bone marrow organoids as a niche for leukemic cells. **a**) Representative confocal images of leukemic blast cells (CFSE, green) and the network (CD31, magenta) within BMOs (DAPI, blue) at 4 h, 8 h, 12 h, 16 h, 20 h, 24 h, 48 h, 72 h, and 96 h after their addition. The maximum projection of z-stack acquisition is shown. Scale bar, 100 μm. **b**) Quantitative analysis of the number of CFSE+ cells per BMO over time. Results of two independent experiments. Symbols indicate mean values, and the error bars show the standard deviations. **c**) Quantification of the CFSE intensity of individual homed LSCs over 96 h. Representative data from one out of three independent experiments. Violin plots represent the smoothened distributions with individually analyzed CFSE+ cells represented as grey dots. Dotted horizontal lines indicate the medians.

## Discussion

Laborious experimental set ups are needed to access living BM; thus it is of high interest to develop *in vitro* assays that provide an easily reachable BM environment for disease modeling. Previous studies have created *in vitro* BM niches to assay HSPC maintenance, expansion, and differentiation using scaffolds, gels, *in vitro-in vivo* hybrid systems, bioreactors, or microfluidic chips^12,20,21,39–44^. Instead, the focus of our work was directed towards the development of a self-organized, highly reproducible BM-mimicking platform, which allows for the investigation of stem cell behavior in a scalable manner. Here, we present a dual-cell type BMO system as a novel approach that recapitulates key aspects of human BM, provides a 3D environment for HSPCs, and mimics the HSPC engraftment dynamics *in vitro.* Furthermore, diseased HSPCs, such as CD34^+^ leukemic blasts, showed a similar migration potential, opening the possibility of performing pharmacokinetic and toxicity studies of novel drugs within such a 3D *in vitro* assay prior to *in vivo* studies. These BMOs were built by co-aggregating human MSCs and ECs, which self-organize into spherical aggregates containing an endothelial network structure. Interestingly, within this 3D co-culture system, a population of MSCs was maintained while other MSCs partially differentiated towards chondrocytes, adipocytes, and osteoblasts, indicated by the deposition of extracellular matrix proteins and specific transcription factors. Calcium deposits were not detected, probably due to the short culture period of these BMOs, considering standard osteogenic differentiation protocols of this process take three or more weeks^45^. The adipo-osteogenic balance of MSCs, which we detected in our system, mimics an essential part of the natural niche environment that is known to be crucial for hematopoiesis^3,46^. Additionally, the deposition of extracellular matrix proteins by MSCs in BMOs could serve as a reservoir for growth factors and provide non-cellular ligands, which is known to be a critical process for the migration and homing of HSPCs in BM^47^. Overall, our results confirmed previously described mesenchymal 3D cultures^19^, including the maintenance of the MSC fate, the deposition of extracellular matrix proteins, and as demonstrated for the first time, the presence of differentiated mesenchymal phenotypes within dual-cell type BMOs.

Importantly, most studies presenting *in vitro* BM systems used scaffolds^18,48^, gels^25,42,49–51^, or microfluidic devices^20,21,41,43,52^ to obtain complex endothelial networks *in vitro.* We demonstrate here that ECs have the potential to self-organize in a complex architectural network within mesenchymal cells without additional support, reminiscent of the organization found in the vascular BM. The intricate balance between the two cell types is highlighted by the finding that elevated numbers of starting ECs did not result in more ECs within BMOs, neither formed a larger network. These results suggest that EC survival and network formation is limited by the secretion of cytokines and/or the disposition of an adhesive substrate by mesenchymal cells in BMOs. Therefore, we hypothesize that a further characterization of the dependencies on various medium components, nutrition availability, or donor origin of the two cell types will ultimately help to refine conditions enabling BMO maintenance for longer times.

Intriguingly, a homing-like behavior of HSPCs was observed when these stem cells were added to fully formed BMOs, and the migration capacity of these stem cells was positively influenced by the abundance of ECs. *In vivo* studies have shown that such cell-cell interactions are essential for the migration, mobilization, and maintenance of HSPCs within BM^23^. Additionally, it is hypothesized that the first interaction during the homing of HSPC in BM is with the sinusoidal endothelium^34^. Therefore, our findings that HSPCs migrated more efficiently in BMOs containing a larger network-like structure, as well as the close proximity between the two cell types, suggest that our 3D system mimics essential parts of such cellular interactions found in natural BM^23,34^. Further investigation of the migration process of HSPCs in BMOs proposed a partial regulation of the CXCL12-CXCR4 signaling pathway, which is known to be the main regulator for human stem cell homing^35^. Only a partial blockage was achieved through the CXCL12-CXCR4 axis, suggesting the involvement of additional signaling pathways. In murine models, evidence was found that CD82, a tetraspanin, is also involved in the regulation of this process^53^. Thus, this BMO system could be used as an *in vitro* tool to investigate such cell-cell interactions and molecular signaling pathways of the human HSPC-niche environment. In this work, we showed that BMOs provide a BM-mimicking environment that recapitulates natural HSPC homing behavior. However, the preservation of the HSPC capacity still needs to be assessed through xenotransplantation assays in future studies^54^.

Most importantly, we found that CD34^+^ leukemic blasts have the potential to efficiently migrate and reside in BMOs, similar to healthy HSPCs. The high expression of CXCR4 on CD34^+^ leukemic blasts suggest that their migration capacity is also regulated over the CXCL12-CXCR4 axis. This hypothesis is supported by the evidence that the CXCL12-CXCR4 signaling pathway is crucial for leukemic blast survival as well as for *in vivo* homing and repopulation of the healthy hematopoietic niche^55,56^. It is known that the vascular density is increased in the AML marrow, while the role of the sinusoidal endothelium on leukemic blasts is still not understood. Thus, our BMOs, which contain a complex endothelial-like network, provide an *in vitro* environment to investigate and understand such endothelial-leukemia cell interactions in future studies. Additionally, this complex *in vitro* BM environment opens up exciting perspectives towards new understandings of clinically relevant mechanisms of resistance as well as of the variability of leukemic blast responses to potential cancer therapeutics.

Overall, we believe that by using the self-assembling potential of MSCs, ECs, and HSPCs/leukemic blasts in 3D, the natural as well as the diseased BM niche is mimicked more precisely than when these interactions are guided in other models (e.g. microfluidic devices). In-depth analysis of the secreted cytokine cocktail derived from dual-cell type BMOs could help to decipher the interplay between the niche and HSPCs/leukemic blasts behavior. Furthermore, the relevance of our system may be enhanced through the addition of even more niche-like cells, such as perivascular^57^, osteoblastic^58,59^, or nerve^60^ cells.

In conclusion, an *in vitro* assay of next-generation BMOs provides an experimental tool for questions related to basic BM niche biology, such as specific cell-cell interactions, homing behavior, or the maintenance of HSPCs and leukemic blasts. Furthermore, the use of BMOs as a testing ground prior to focused *in vivo* experimentation could reduce the number of experimental animals, and thereby, reduce overall costs. The advantages of this system over standard *in vitro* assays are *i)* the presence of multiple BM cell types, which creates an *in vivo*-mimicking environment for HSPCs and leukemic blasts, *ii)* the 3D context, allowing for systematic searches for dosing regimens, which could grant better pre-screening data for future *in vivo* studies, and *iii)* the potential to investigate patient-to-patient variability in terms of drug resistance. We additionally believe that the reproducibility and scalability of the Gri3D microwell platform meets crucial requirements for various applications of BMOs from large-scale compound screenings to patient-specific tests^61^. In summary, the multi-cell type 3D BMOs represent an innovative preclinical model for recapitulating hematopoietic and leukemic stem cell dynamics in a complex BM-like niche environment.

## Methods

### Cell sources

Human samples were received from consenting donors from the University Hospital Zurich and the Geneva University Hospital in compliance with Swiss law. Frozen vials of MSCs were purchased from Lonza or received from the University Hospital Zurich. HUVECs were received as frozen vials from the University Hospital Zurich. Umbilical cord blood (UCB) was collected from consenting donors from the University Hospital Zurich. BM or peripheral blood (PB) samples from AML patients at diagnosis were obtained with informed consent and approved by the local ethics committee (Medical Ethics Committee of the Canton Geneva; Study no. 2020-00176). The diagnosis and classification of leukemic samples was performed according to the WHO 2016 classification. The mononuclear fraction of CB and AML cells were isolated by using a Ficoll density centrifugation with Lymphoprep (Axonlab, 1114545) in Leucosep tubes (Greiner bio-one, 227290). Prior to culturing or freezing AML cells, the mononuclear fraction was further enriched for CD34^+^ cells using the UltraPure CD34 Microbead kit (Miltenyi Biotec, Bergisch Gladbach, Germany, 130-100-453) and magnetic-activated cell-sorting separation columns (Miltenyi Biotec, 130-042-201). The mononuclear CB samples were cryopreserved with serum-free freezing medium (FILOCETH^62^) and stored in liquid nitrogen.

### Magnetic depletion and enrichment of cord blood and AML cells

The cryopreserved CB vials were thawed and diluted dropwise with IMDM plus GlutaMAX (1x, Gibco, 31980-030) containing 0.5 % BSA Fraction V (Gibco, 15260-037) and 10 μg/ml DNase I (Roche, 11284932001). The cryopreserved AML cells were thawed by adding dropwise StemSpan SFEM II medium (StemCell Technologies, 09605) supplemented with 1 % penicillin/streptomycin (P/S) (Gibco, 15140-122). The Dead Cell Removal Kit (Miltenyi Biotec, 130-090-101) was used to remove apoptotic and dead cells via magnetic depletion according to the indicated instructions. The CB cells were lineage depleted with the human lineage cell depletion kit (Miltenyi Biotec, 130-092-211), which contains the following biotin-conjugated monoclonal antibodies, CD2, CD3, CD11b, CD14, CD15, CD16, CD19, CD56, CD123, and CD235a (Glycophorin A). The Lin^−^ fraction of the CB cells and the AML cells were further enriched for CD34 with the human CD34 MicroBead Kit UltraPure (Miltenyi Biotec, 130-100-453), which contains microbeads conjugated to monoclonal mouse anti-human CD34 antibodies (isotype: mouse IgG1). For the depletion or enrichment, the cells were resuspended in phenol red-free IMDM (1x, Gibco, 21056-023) or PBS (pH 7.4, 1x, Gibco, 10010-015) containing 0.5 % BSA Fraction V, 2 mM EDTA (pH 8.0, Gibco, 15575-020), 1 % P/S and optionally 20 mM Hepes (Gibco,15630-056). The enrichment or depletion was performed according to the manufacturer’s protocol and run over the LS or MS columns (Miltenyi Biotec, 130-042-401 or 130-042-201, respectively) depending on the total cell number.

### Flow Cytometry

The purity check of the lineage depletion or enrichment of HSPCs was performed by staining with the following monoclonal directly conjugated antibodies CD45-PECy5 or CD45-AF700, CD34-BV421 or CD34-PE, CD38-APC and optionally with the biotin lineage depletion cocktail and streptavidin-AF488 (**Table 1**). The CD34^+^ AML cell purity were assessed by staining cells with anti-human CD34-PE-Vio770 and CD45-VioGreen (**Table 1**). To assess the expression levels of CXCR4, the CD184-BV605 monoclonal antibody was used (**Table 1**). The single cells from the digested BMOs were stained and analyzed with the following monoclonal directly conjugated antibodies: CD71-AF700, CD31-APCeFluor780 or AF700, CD105(ENG)-FITC, CD146-BV711, VEGFR-PE, CD144-PECy7, CD45-PECy5, CD34-BV421, and CD38-APC (**Table 1**).

**Table 1.**

| Antibody | Fluorophore | Reactivity | Host/Isotype | Company | Reference | Clone | Dilution |
| --- | --- | --- | --- | --- | --- | --- | --- |
| CD71 | AF700 | Human | Mouse IgG2a κ | BD Pharmingen | 563769 | M-A712 | 1:40 |
| CD31 | AF700 | Human | Mouse IgG1 κ | Biolegend | 3031134 | WM-59 | 1:320 |
| CD31 | APC eFluor780 | Human | Mouse IgG1 κ | eBioscience | 47-0319-41 | WM-59 | 1:320 |
| ENG | FITC | Human | Mouse IgG1 κ | BD Pharmingen | 561443 | 266 | 1:80 |
| CD146 | BV711 | Human | Mouse IgG1 κ | BD Pharmingen | 563186 | P1H12 | 1:160 |
| VEGFR | PE | Human | Mouse IgG1 κ | R&D | FAB357P-025 | 89106 | 1:160 |
| CD144 | PECy7 | Human | Mouse IgG1 κ | Invitrogen | 25-1449-42 | 16B1 | 1:640 |
| CD45 | PECy5 | Human | Mouse IgG1 κ | BD Pharmingen | 560974 | HI30 | 1:320 |
| CD45 | AF700 | Human | Mouse IgG1 κ | BD Pharmingen | A79390 | J33 | 1:40 |
| CD45 | VioGreen | Human | Cell line IgG1 | Miltenyi Biotec | 130-110-638 | REA747 | 1:50 |
| CD34 | BV421 | Human | Mouse IgG1 κ | BD Pharmingen | 562577 | 581 | 1:160 |
| CD34 | PE | Human | Mouse IgG1 κ | BD Pharmingen | 555822 | 581 | 1:10 |
| CD34 | PE-Vio770 | Human | Mouse IgG1a κ | Miltenyi Biotec | 130-113-180 | AC136 | 1:50 |
| CD38 | APC | Human | Mouse IgG1 κ | BD Pharmingen | 555462 | HIT2 | 1:10 |
| CXCR4 | BV605 | Human | Mouse IgG2a κ | Biolegend | 306521 | 12G5 | 1:40 |
| Streptavidin | AF488 | Biotin | - | Life Technologies | S32354 | - | 1:1000 |

For live/dead cell discrimination, 4′,6-diamidino-2-phenylindole (DAPI, Sigma, D9542), propidium iodide (PI, Sigma, 81845), or 7-Amino-Actinomycin D (7-AAD, Beckman Coulter, B88526) was added to the cell suspensions. UltraComp eBeads (Invitrogen, 01-2222-42) were used to create single color controls compensation. The LSRII, LSR Fortessa (Becton Dickinson) or Navios 10-color (Beckman Coulter) flow cytometers were used to analyze the samples. Data was analyzed in Flowjo (Version 10.5.3) or Kaluza (Analysis Version 2.1) and fluorescence minus one controls (FMOs) were used to set the gates (data included in Supp. Figures).

### 2D Cell Culture

Human MSCs were expanded on 2D flasks in αMEM (Gibco, 12571-063) supplemented with 10 % fetal bovine serum (FBS, Gibco, 10500-064), 2 mM GlutaMax, 1 % P/S and 1 ng/ml human FGF2 (Peprotech, 100-18B). When the MSCs reached 80-90 % confluency, the cells were detached with 0.25 % Trypsin (Gibco, 25200-072) and partially re-plated in expansion medium or used for the 3D culture. For cryopreservation the MSCs were stored in 90 % FBS containing 10 % DMSO (Sigma, D2438).

HUVECs were expanded on 2D flasks in EGM-2 (Lonza, CC-3162) or ECGM (Provitro 211 1101). If the confluency reached around 80-90 % the HUVECs were detached with TrypLE™ Express (Gibco, 12605-028) and partially re-plated in expansion medium or used for the 3D culture. The cryopreservation was done in a medium containing 60 % EGM-2 or ECGM, 30 % FBS and 10 % DMSO.

CD34^+^ leukemic blasts were cultured in StemSpan™ SFEM II serum-free medium (STEMCELL Technologies, 09605) supplemented with the StemSpan™ CD34^+^ Expansion supplement (STEMCELL Technologies, 02691) and 1 % P/S. Cells were initially plated in 96-well culture plates at a concentration in the range of 5-10 × 10^4^ cells/mL and incubated at 37 °C in a humidified atmosphere of 5 % CO_2_ in air. Complete medium change was performed every third day. Cultures were maintained for several months. All cells were cryopreserved in liquid nitrogen until usage.

### 3D Cell Culture

For the 3D culture of MSCs and HUVECs, cells were seeded in different ratios in the Gri3D cell culture platform (SUN bioscience, Gri3D-24-S-24P, Gri3D-96-S-24P)^22^. The 24 or 96 well plate format of the Grid3D platform was used. Each well contained an array with 121 PEG μ-wells with a diameter of 400 μm. For the control condition (100/0) 500 MSCs were seeded per μ-well. For the coaggregation of MSCs and HUVECs, either 250 MSCs and 250 HUVECs (50/50) or 375 MSCs and 125 HUVECs (75/25) were seeded per μ-well. The cells were cultured in previously published medium^19,24^ that contains the following components: DMEM/F12 (Gibco, 31331-028) and human endothelial SFM (Gibco, 11111-044) supplemented with 0.1 mM 2-mercaptoethanol (50 mM, Gibco, 31350-010), 1x non-essential amino acids (Gibco, 11140-035), 1x N2 (Gibco, 17504-001), 1x B27 supplements (Gibco, 17504001), 1 % P/S, recombinant human fibroblast growth factor 2 (hFGF-2, Peprotech, 100-18B), recombinant human epidermal growth factor (hEGF, 20 ng/ml, Chimerigen Laboratories, CHI-HF-210EGF), recombinant human platelet derived growth factor (hPDGF-AB, 20 ng/ml, Merck, GF106), recombinant human oncostatin M (hOSM, 20 ng/ml, Gibco, PHC5015) and recombinant human insulin-like growth factor 1 (hIGF-1, 40 ng/ml; Peprotech, 100-11) and 15 % chick embryo extract (CEE)^19,24^. CEE was produced according to published protocols^63,64^. The day of seeding MSCs and HUVECs in μ-wells is referred to as Day 0. The full medium was replaced every second or third day. The addition of the CD34^+^ HSPCs was performed on day 3 when compact BMO were typically formed. 62.5 to 750 human lineage-depleted (Lin^−^) CB cells, CD34-enriched (CD34^+^) HSPCs or CD34^+^ AML blasts were seeded per BMO on day 3. The CD34^+^ HSPCs and CD34^+^ AML blasts were labelled with the CFSE cell division assay kit (Cayman, 10009853), according to the manufacturer’s instruction prior to their seeding on BMOs. Every 4h for the initial 24h and then after 48h, 72h and 96h BMO were collected for further analysis. If the CD34^+^ cells were added to the BMOs, the collection was performed using a FACS strainer (Falcon, 352235) to remove non-homed cells. For the inhibition of the homing potential, the CD34^+^ HSPCs were treated with 100 μM of AMD3100 octahydrochloride (Tocris, 3299) or with 8 μM of SB290157 (Sigma, SML1192) for 45 min at 37 °C. The anti-PTN antibody (R&D, AF-252) was added in a 1/25 dilution to the culture medium of the BMOs for 45 min at 37 °C. For the cytometric analysis of the surface markers the BMOs were digested with PBS containing 3 mg/ml collagen IV (Gibco, 17104-019), 4 mg/ml Dispase (Gibco, 17105-041) and 2 mg/ml DNase I (Roche, 11284932001) to obtain a single cell suspension.

### Immunostaining

The collected BMOs were fixed with 2 % paraformaldehyde (PFA, ThermoFischer Scientific, 15434389) for 1 h and permeabilized with 0.3 % Triton-X-100 (Sigma, T8787) in PBS for 3 h at RT. Depending on the secondary antibody either 1 % Albumin Fraction V (AppliChem, A1391.0500) or 10 % donkey-serum in PBS 0.01 % Triton-X-100 as blocking solution was used at 4 °C overnight. After the blocking step, the following primary antibodies were used for the immunostainings (**Table 2**) in combination with the below listed secondary antibodies (**Table 3**). DAPI was used to mark the nuclei.

**Table 2.**

| Antibody | Species Reactivity | Host/Isotype | Class | Company | Reference | Dilution |
| --- | --- | --- | --- | --- | --- | --- |
| CD45 | Human | Rabbit | Monoclonal | Abcam | Ab40763 | 1:200 |
| Endoglin | Human | Goat | Polyclonal | R&D | AF1097 | 1:100 |
| Nestin | Human | Chicken | Polyclonal | Abcam | Ab81755 | 1:100 |
| CD31 | Human | Mouse | Monoclonal | Cell Signaling | 89C2 | 1:200 |
| CD144 | Human | Rabbit | Polyclonal | Abcam | Ab33168 | 1:200 |
| TAGLN | Human | Rabbit | Polyclonal | Abcam | Ab14106 | 1:200 |
| RUNX2 | Human | Mouse | Monoclonal | Abcam | Ab76956 | 1:100 |
| Sox9 | Human | Rabbit | Monoclonal | Abcam | Ab185966 | 1:100 |
| PTN | Human | Goat | Polyclonal | R&D | AF-252 | 1:50 |
| CD34 | Human | Rabbit | Monoclonal | Abcam | Ab81289 | 1:100 |

**Table 3.**

| Target | Host | Conjugation | Company | Reference | Dilution |
| --- | --- | --- | --- | --- | --- |
| Rabbit | Donkey | A488 | ThermoFisher Scientific | A-21206 | 1:500 |
| Rabbit | Donkey | A568 | ThermoFisher Scientific | A-10042 | 1:500 |
| Rabbit | Donkey | A647 | ThermoFisher Scientific | A-31573 | 1:500 |
| Goat | Donkey | A568 | ThermoFisher Scientific | A-11057 | 1:500 |
| Mouse | Donkey | A647 | ThermoFisher Scientific | A-31571 | 1:500 |
| Chicken | Donkey | A488 | Jackson ImmunoResearch | 703-545-155 | 1:500 |
| Rat | Donkey | A488 | ThermoFisher Scientific | A-21208 | 1:500 |

### Microscopy

For the growth of the BMOs, brightfield images of the Grid3D array were acquired every day or every second day on a Nikon Ti Microscope with a 4x objective (N.A. = 0.13, air) and automated stitching of tiles to form large images. Confocal images for qualitative representation were acquired on an inverted LSM700 microscope (Zeiss) with a 20x objective (N.A. = 0.8, air). The pixel size was set to 0.16 μm and a z-step of 3 μm, pinhole set a 36.8 micron (equivalent DAPI: 1.28 A.U. = 1.8 μm, A647: 0.91 A.U. = 2.3 μm, A488: 1.13 A.U. = 2.2 μm, A568: 1.09 A.U. = 2.2 μm) was used. Confocal images, which were analyzed with a custom script (see methods: Image analysis), were acquired on an upright LSM710 microscope (Zeiss) or an upright SP8 microscope (Leica), with 20x objectives (N.A. = 0.8, air and N.A. = 0.75, air, respectively). For both, the pixel size was set to 0.71 μm and a z-step of μm was used. For the LSM710 the pinhole was set to 30.4 microns (equivalent DAPI & A647: 1.15 A.U. = 1.6 μm; A488: 0.98 A.U. = 1.7 μm; A568: 0.86 A.U. = 1.7 μm) and for the SP8 the pinhole was set to 47.5 microns (equivalent DAPI: 0.87 A.U. = 0.7 μm; A488: 0.75 A.U. = 0.8 μm; A568: 0.65 A.U. = 0.8 μm; A568: 0.58 A.U. = 0.9 μm). The histology sections were acquired with an upright DM5500B (Leica). The color camera CMOS DMC 2900 was used to acquire the images with a 10x (N.A. = 0.3, air), 20x (N.A. = 0.7, air) or 40x (N.A. = 0.5-1.0, oil) objective.

### Image analysis

Growth of BMOs was analyzed in Fiji (2.0.0-rc-69/1.52n) with a custom script, which automatically detects the BMO area by thresholding and particle analysis within the array on brightfield images. The Matlab EasyXT-User Interface [1] for Imaris was used to detect the Spheroid and the Network as surface objects, and to detect the CD45-positive or CFSE-traced cells as spots or surfaces objects. The whole organoid surface was detected using smoothened ENG or DAPI signal and the vascularization network based on the CD31 staining. Mask channel for the individual detected surfaces were generated for further analysis. For the network architecture, a FIJI [2] custom script was used to extract the vascular network mask from the in Imaris image, skeletonized the data and analyzed the skeleton^65^. The script relies on the « BIOP Basics » [3] ActionBar to facilitate processing.

1. EasyXT (https://github.com/BIOP/EasyXT) and EasyXT-GUI (https://github.com/BIOP/EasyXT-GUI)
2. FIJI (https://www.fiji.sc)
3. BIOP Basics (https://c4science.ch/w/bioimaging_and_optics_platform_biop/image-processing/imagej_tools/biop-basics/)

For the estimated BMO volume, a FIJI custom script that extracts the BMO mask made in Imaris was used, makes a z-projection and runs “Analyze particles” with the “fit ellipse” option. From the measurements, it gets the major and minor radii and calculates an estimation of the volume with the formula: 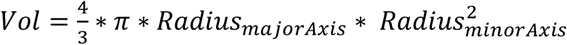. Customized analysis was performed to measure the distances from the cells to the organoid surface and to the vascularization, respectively. Using EasyXT in Matlab, new channels were generated in Imaris whose voxel’s intensities equal the 3D-distance towards the above rendered organoid and vascular network surfaces, respectively. For each detected cell, the distance towards the organoid or the network surface was approximated by the average intensity of the generated distance channels within the cell volume. To give a reference, the expected average distances were modeled per organoid by averaging the intensities of the distance channels within the whole organoid volume (for the surface-distance modeling) or the network volume-cleared organoid volume (for the network-distance modeling).

### Histology

BMOs were collected from the array as described above and fixed with 2 % or 4 % PFA or 10 % Formalin (Sigma, HT501128) overnight at 4 °C. For paraffin sectioning, the BMOs were embedded in a drop of HistoGel (ThermoFisher Scientific, HG-4000-012). The dehydration of the hardened Histogel drops was done in the automated dehydration machine (Sakura, VIP6). Afterwards, the samples were embedded in paraffin (Merck, 1.11609.2504) according to standard histological procedures and 4 μm thick sections of the paraffin blocks were cut on the microtome (Leica, HM325). The paraffin sections were stained with Haematoxylin-Eosin (HE) to assess general morphology. Alcian blue (AB) at pH 2.5 was used to stain mucus, Alizarin Red (AR) for calcium deposits, Periodic Acid Schiff (PAS) for polysaccharides, Sirius Red (SR) stains collaged, Marius Scarlett Blue (MSB) is for fibrin and Miller (M) stains for elastin.

For the cryosections, the BMOs were placed in an increasing sucrose solution (7.5 % for 1 h, 15 % for 1 h and 30 % O/N; Sigma, S1888) for improved cryoprotection. Afterwards, the BMOs were embedded in a drop of 7.5 % gelatin solution (Sigma, G1890) with phosphate buffer (Sigma, 71505 and S7907) 0.12 M and 15 % sucrose and placed on an organoid embedding sheet (StemCell Technologies, 68579A). The drop of gelatin was left to harden at 4 °C for 15 min. The gelatin drop was placed in a mold half-filled with gelatin (Sigma, G2500) or cryomatrix (Thermo Scientific, 6769006). The block was slowly frozen in isopentane (VWR, 24872.298) cooled down in a bath of ethanol (Fischer Chemical, E/0600DF/15) and dry ice to −65 °C to - 70 °C. The cryoblocks were cut into 8 μm thick sections on the cryostat (Leica, 1850 or 1950). Oil Red O staining was used to specifically highlight lipids.

### Statistical Analysis

The according statistical analysis is indicated in each figure’s description. Student’s t-tests were used to compare two data sets, while ordinary one-way ANOVA with multiple comparisons was used for a group of data. All data sets are represented with mean, median and standard deviations or median quartiles as indicated. Statistical significance was set at a p-level of less than 0.05.

## Acknowledgments

We thank Stefano Davide Vianello for the design of the schematic BMO layouts and various feedback on the manuscript and François Rivest for the creation of the automated script for the detection of the BMO area. We thank the staff of the bioimaging and optics platform (BIOP) for their support and advice regarding image acquisition, processing and quantification, Jessica Sordet-Dessimoz and all the members of the histology core facility (HCF) for their help with histological stainings and sample processing as well as interpretation of the results, and the flow cytometry core facility (FCCF) members for their support with the flow cytometry experiments.

## Author Contributions

M.P.L., S.G. and S.H. conceived the study. S.G., M.H. and M.M-L. performed experimental work and M.H. and S.G. did the statistical analyses; S.G., M.H. and M.P.L. interpreted results and wrote the manuscript; S.H. contributed to the experimental work by the establishment of the mouse system; S.H and N.B. provided the microwell platform Gri3D; M.H. and R.G developed the automated image analysis; M.E., E.K. and K.G provided CB, MSCs and HUVECs samples, M.M-L. and T.M. provided AML samples and supported us with experimental work and know-how.

## Supplementary Figures

**Supp. Fig. 1.**
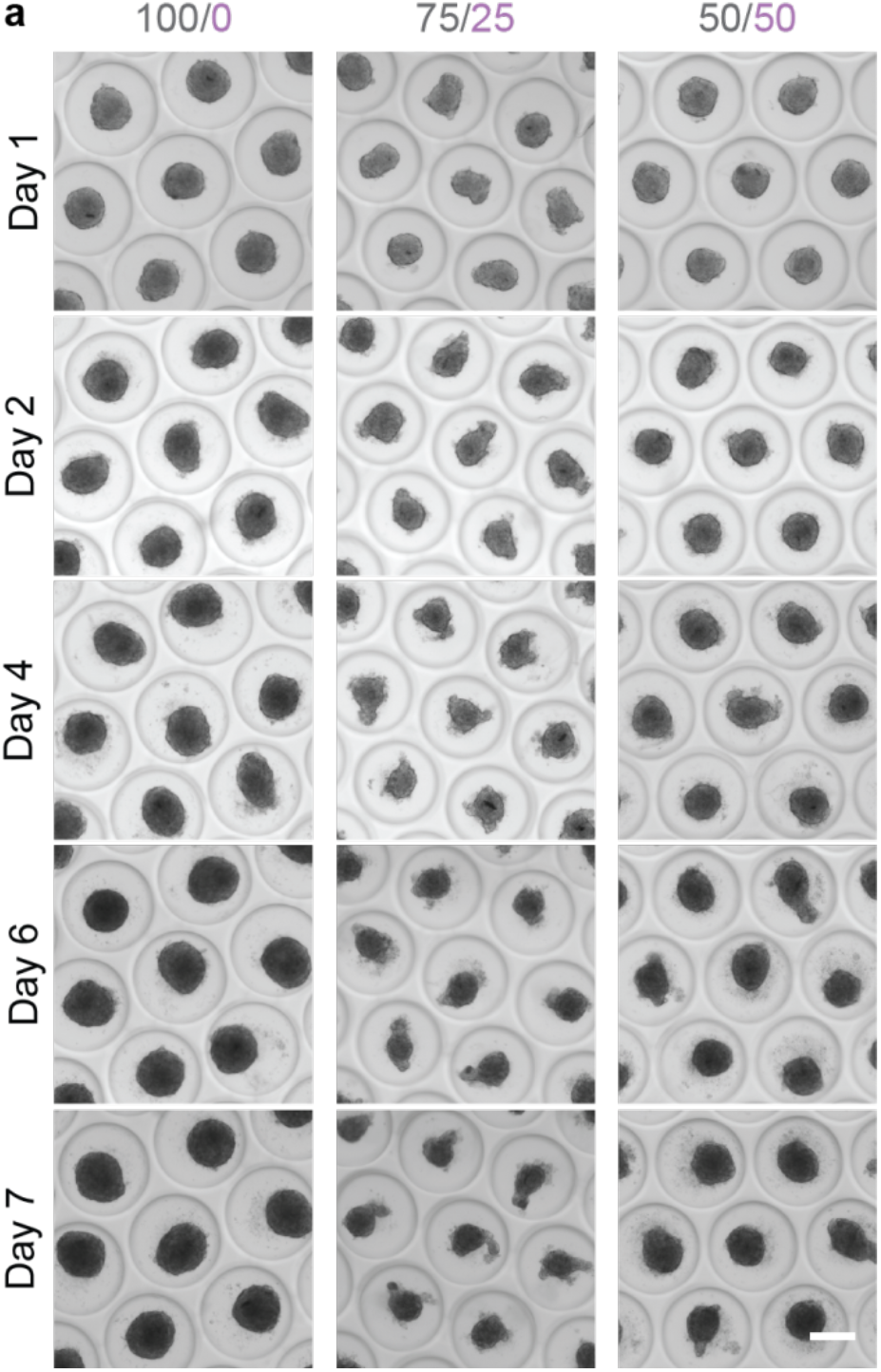
Formation of bone marrow organoids in a scalable manner. Representative brightfield images of the resulting BMOs of aggregation of cells in different ratios (grey: MSCs, magenta: ECs) in the Gri3D platform at day 1, 2, 4, 6, 7. Scale bar, 200 μm.

**Supp. Fig. 2.**
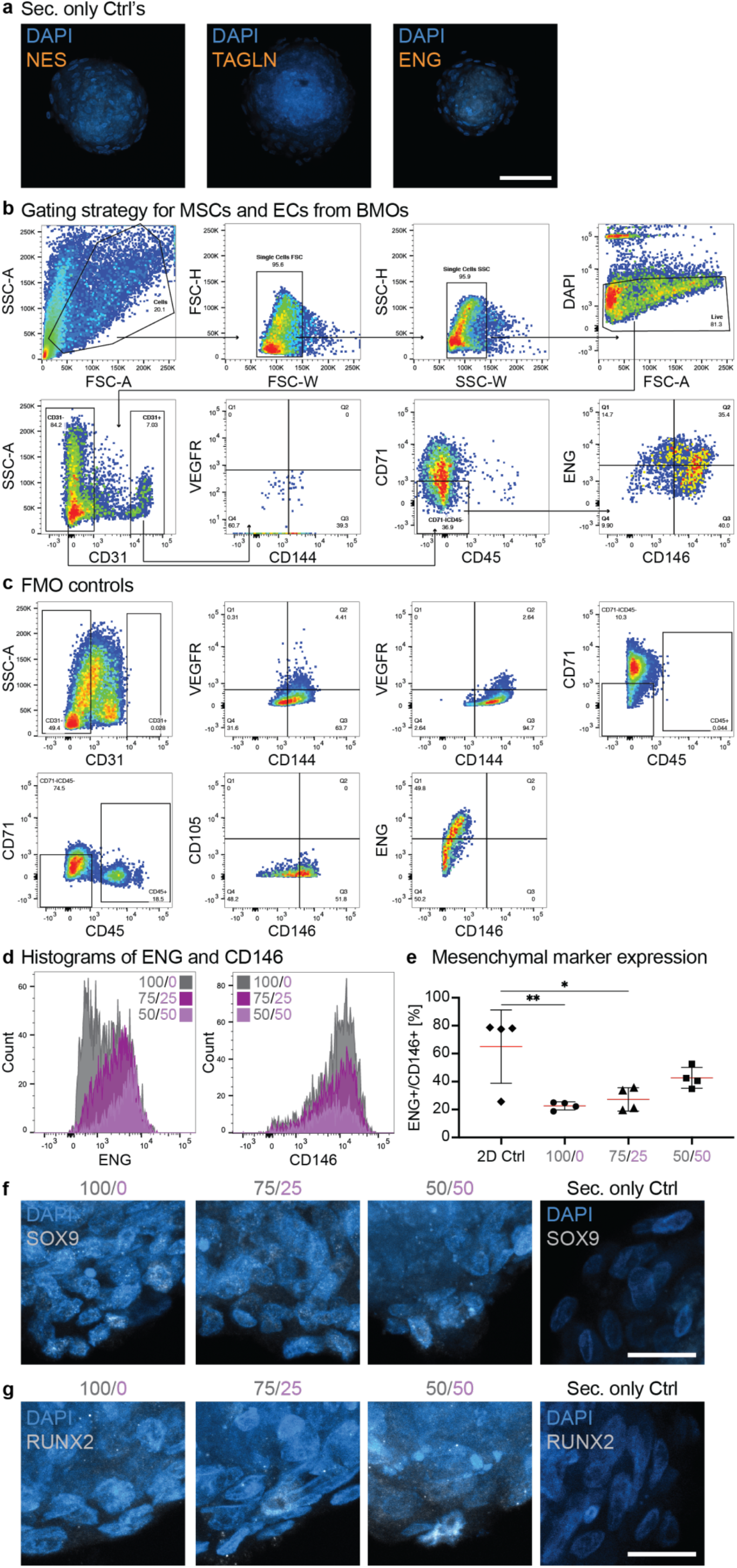
Self-renewing and differentiated MSCs in bone marrow organoids. **a**) The control staining for the immunofluorescent images was done by using only the appropriate secondary antibodies. Images showing the secondary antibody for NES, TAGLN, and ENG. Scale bar, 100 μm. **b**) The gaiting strategy for MSCs and ECs from BMOs to analyze these compartments by flow cytometry. Single cell exclusion is done on forward scatter (FSC) as well as side scatter (SSC). DAPI negative cells mark the live cell population. The live cells are gated for the CD31^+/−^ cell population. The CD31+ population stains the endothelial compartment of BMOs and is further discriminated by the endothelial markers CD144 and VEGFR. The CD31^−^ population is negatively selected for CD71 and CD45; the resulting CD71^−^/CD31^−^/CD45^−^ population marks the mesenchymal compartment of BMOs. Within this CD71^−^/CD31^−^/CD45^−^ population the ENG^+^/CD146^+^ double positive phenotype is investigated. **c**) Fluorescence minus one (FMO) controls of each marker was used to set the appropriate gate. **d**) Representative histogram of the ENG and CD146 expression within the mesenchymal CD71^−^/CD31^−^/CD45^−^ population. The mean fluorescent intensity (MFI) of ENG is 1’671 in the 100/0, 2’408 in the 75/25, and 3’787 in the 50/50 condition. While the MFI of ENG is 7’438 in the 100/0, 7’041 in the 75/25, and 6’109 in the 50/50 condition. **e**) Quantitative analysis of the percentage of CD71^−^/CD31^−^/CD45^−^/CD105^+^/CD146^+^ expression. Red horizontal lines indicate the mean values and the error bars show the standard deviations. Results of four independent experiments. Statistical analysis by ordinary one-way ANOVA and multiple comparisons. *p<0.05; **p<0.01 **f**) Representative confocal images of SOX9 (white) expression in BMOs in all three conditions (100/0, 75/25, 50/50) and the secondary antibody control image for SOX9. **g**) Representative confocal images of RUNX2 (white) expression in BMOs in all three conditions (100/0, 75/25, 50/50) and the secondary antibody control image for RUNX2. **f, g**) Maximum intensity projection of z-stack images. Scale bars, 100 μm.

**Supp. Fig. 3.**
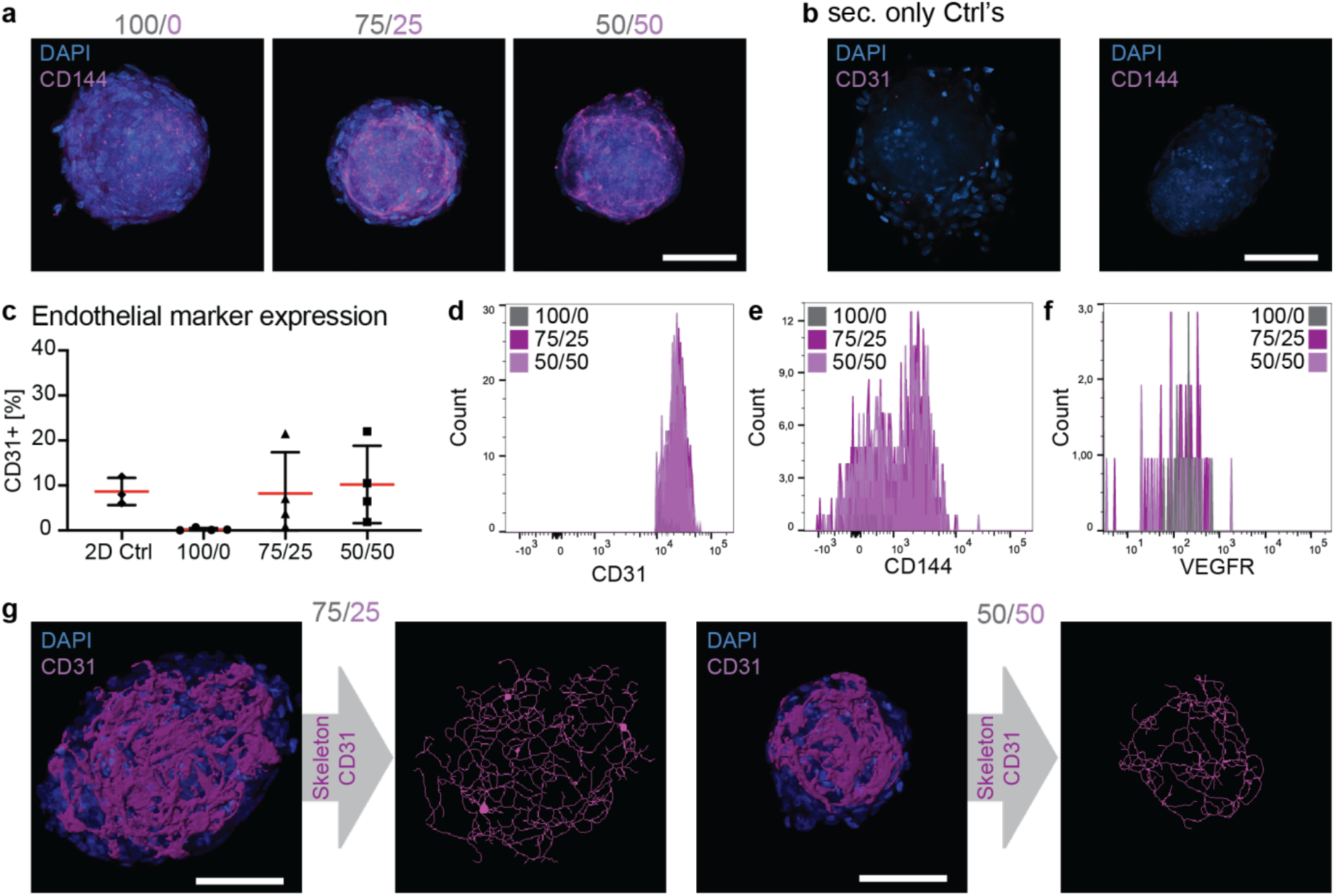
Establishment of a self-organized, vascular-like network. **a**) Representative confocal images of the endothelial marker CD144 (magenta) from BMOs of the three conditions (100/0, 75/25, 50/50). Maximum intensity projection of z-stack images. Scale bar, 100 μm. **b**) Images showing the secondary antibody control for CD31 (magenta) and CD144. Scale bar, 100 μm. **c**) Quantitative analysis of the percentage of CD31^+^ marker expression of live BMO cells. Red horizontal lines indicate the mean values and the error bars show the standard deviations. Results of four independent experiments. Statistical analysis by ordinary one-way ANOVA and multiple comparisons. **d, e, f**) Representative histogram of the CD31, CD144, VEGFR expression of live BMO cells from the three different conditions (100/0, 75/25, 50/50). **g**) Representative 3D rendered data of the CD31 marker and the resulting skeletonized images, which were used for the quantification of network architecture in BMOs of the two multicellular conditions 75/25 and 50/50. Scale bars, 100 μm.

**Supp. Fig. 4.**
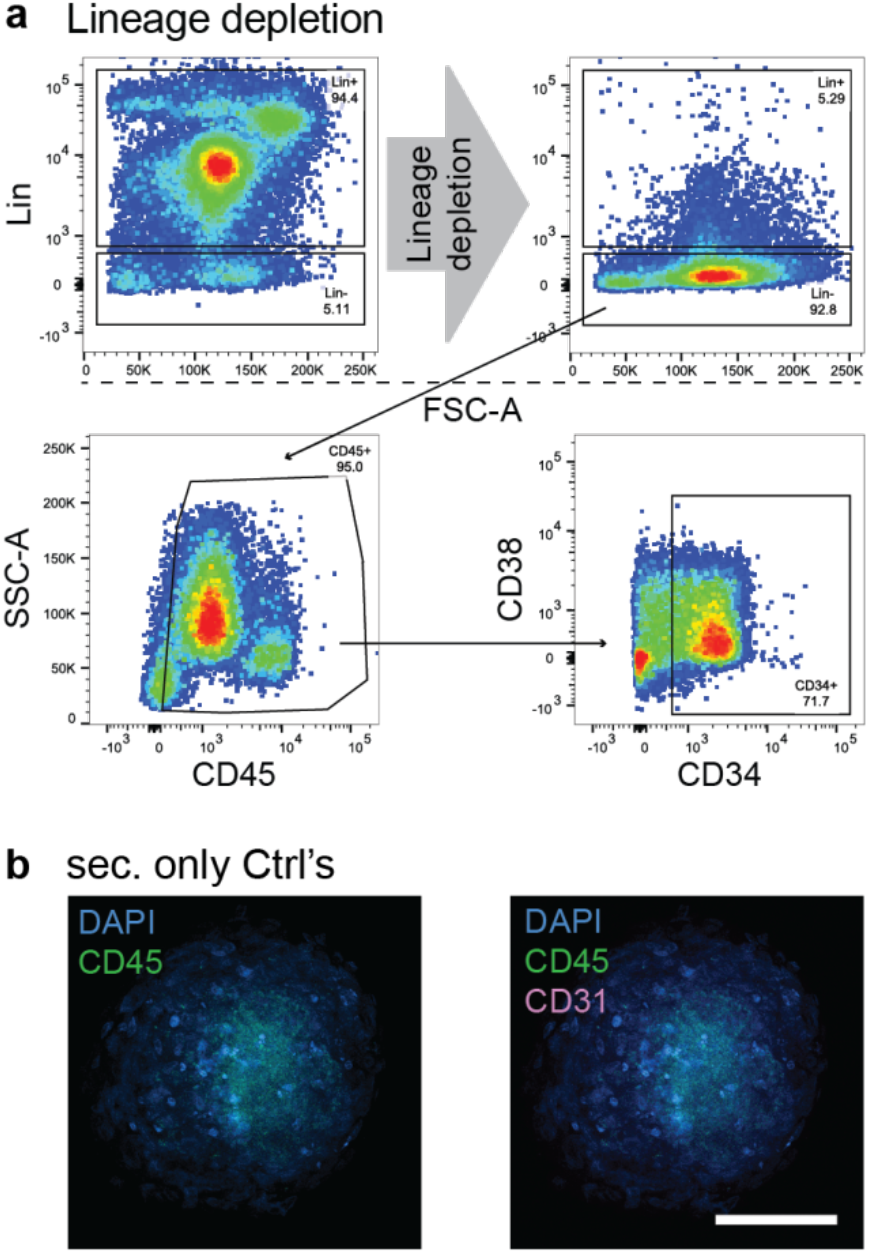
Bone marrow organoids as a 3D migration assay. **a**) Representative flow cytometry plots showing the purity check of the lineage depletion procedure to gain an enriched HSPCs population, which was used for these experiments. **b**) Images showing the secondary antibody control for CD45 (green) and CD31 (magenta). Scale bar, 100 μm.

**Supp. Fig. 5.**
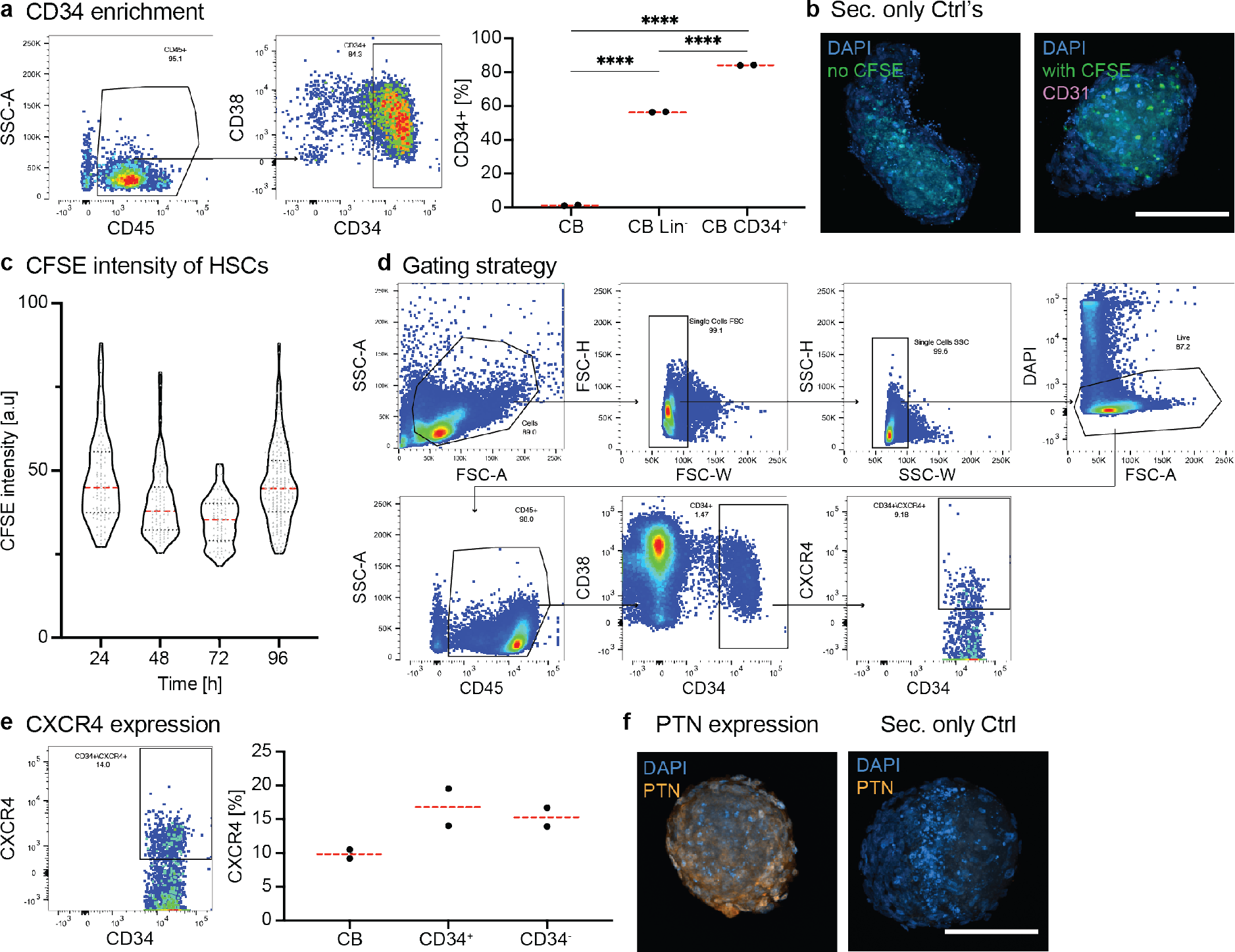
Dynamics and mechanism of CD34^+^ HSPC homing. **a**) Representative flow cytometry plots showing the CD34 enrichment (center panel) of the lineage depleted HSPCs population (left panel). Quantification of the percentage of CD34^+^ cells in the total cord-blood sample compared to the lineage depleted HSPCs, as well as the additional CD34 enriched population (right panel). Statistical analysis by ordinary one-way ANOVA and multiple comparisons. ****p<0.0001. **b**) Images showing the secondary antibody controls. Only DAPI (blue) staining without CFSE^+^ cells (left panel) and DAPI staining with CFSE^+^ cells and only secondary antibody for CD31 (right panel). **c**) Quantification of the CFSE intensity of individual homed CD34^+^ HSPCs over 96 h. Representative data from one experiment out of three independent experiments. Violin plots represent the smoothened distributions with individual analyzed CFSE^+^ cells represented as grey dots. Red dotted horizontal lines indicate the medians and black dotted lines the quartiles. **d**) Gating strategy for the purity check of the depletion and enrichment as well as the CXCR4 gating. **e**) Representative flow cytometry plot of the CXCR4 expression on CD34^+^ HSPCs (left panel). Quantitative analysis of the percentage of CXCR4 in CB, CD34^+^, and CD34^−^ HSPCs samples (right panel). Data from two experiments. Statistical analysis by ordinary one-way ANOVA and multiple comparisons. **f**) Representative confocal images of the PTN (orange) expression in BMOs of the condition 75/25 MSC/EC ratio and the according secondary antibody control for PTN. Maximum intensity projection of z-stack images. Scale bar, 100 μm.

**Supp. Fig. 6.**
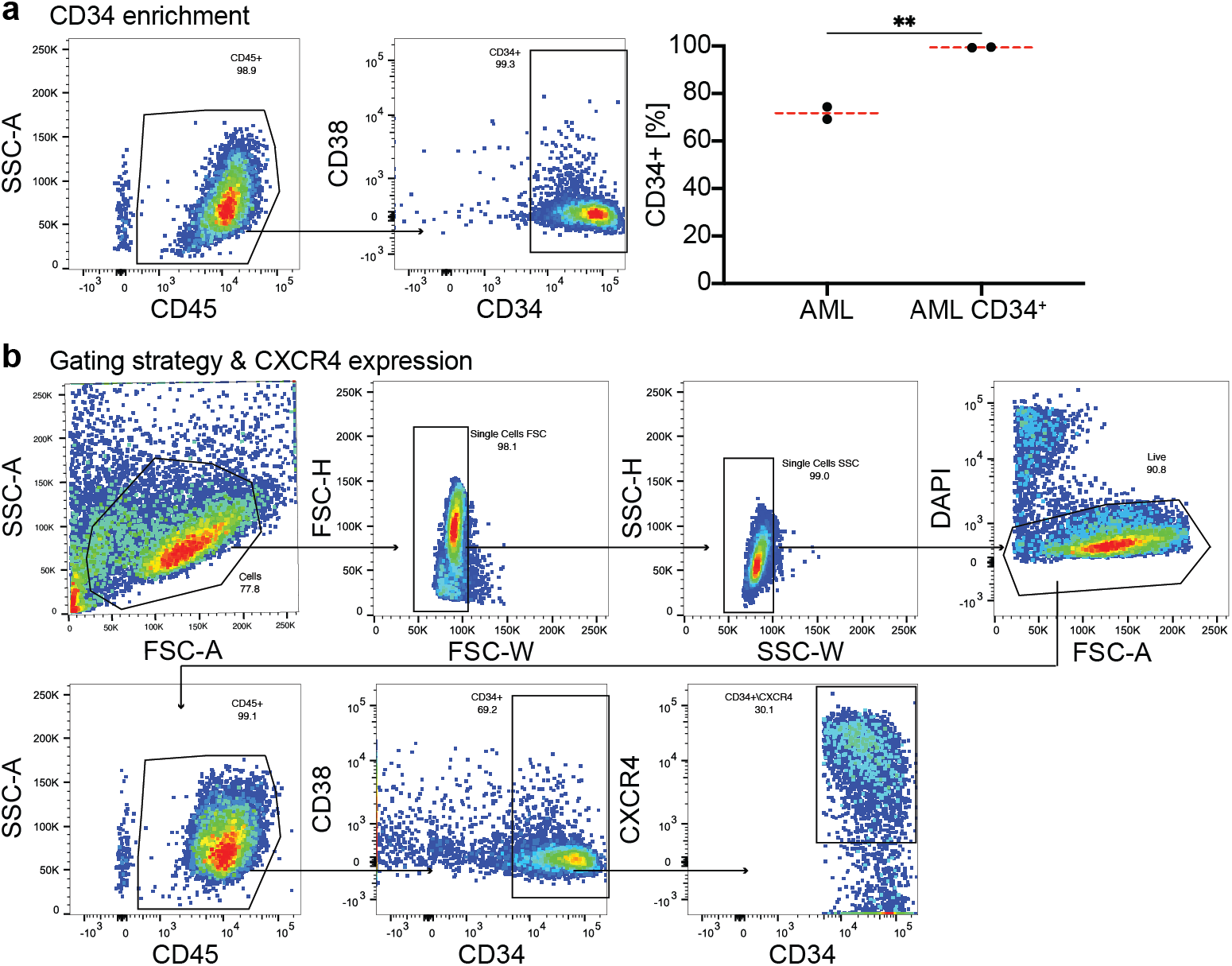
Bone marrow organoids as a niche for leukemic blasts. **a**) Representative flow cytometry plots showing the CD34 enrichment from the AML population (left two panels). The quantification of the percentage of CD34^+^ cells in enriched population compared to the starting AML population (right panel). Data from two independent experiments. Statistical analysis by Student’s t-test. **p<0.01. **b**) Gating strategy for the purity check of the depletion and enrichment as well as the CXCR4 gating on a representative AML sample.

## Notes

### Competing Interest Statement

The authors have declared no competing interest.

